# Exitron-type splicing of intrinsically disordered regions recurs across jawed vertebrates despite sequence divergence: donor-proximal *cis* control in *TARDBP*

**DOI:** 10.64898/2026.10.06.756961

**Authors:** Takuma Yamagishi, Shingo Koide, Genri Toyama, Aya Washida, Shunta Okawara, Kohta Yoshida, Osamu Onodera, Akihiro Sugai

**Author notes:** Corresponding author: Akihiro Sugai.

## Abstract

Intrinsically disordered regions (IDRs) mediate molecular interactions but can also promote aberrant self-association. Using donors embedded within IDR-coding sequences, exitron-type splicing can remove these sequences, giving them both protein-coding and splice-regulatory roles. Yet IDR amino-acid sequences often diverge rapidly, raising the question of whether and how their coding sequences support IDR-removing splicing across species. Using RNA-seq data from jawed vertebrates, we identified events predicted to remove most of an IDR while preserving annotated structured regions. Across species, donor-flanking nucleotide identity exceeded expectations from synonymous alternatives encoding the same amino-acid sequence. *TARDBP*, which encodes TDP-43, provided a model for examining how coding sequences both encode IDRs and regulate exitron-type splicing. Sequence-divergent vertebrate *TARDBP* terminal regions supported this splicing in human cells. In teleosts, *TARDBP* ohnologs (*tardbpa* and *tardbpb*) encoded compositionally similar disordered tails but differed markedly in IDR-removing exitron-type splicing. Reciprocal donor-proximal substitutions in zebrafish minigenes redirected donor choice and altered the balance between IDR-retaining and IDR-removing transcripts. In corresponding reporters, these substitutions altered protein-product balance and cytoplasmic puncta formation. These findings show that IDR-removing exitron-type splicing can recur despite sequence divergence and that donor-proximal *TARDBP* sequences both encode IDR segments and regulate IDR inclusion, linking RNA processing to protein behavior.

## Introduction

Protein-coding sequences can simultaneously encode amino acids and harbor RNA-regulatory information. Exonic splicing enhancers illustrate this dual use: by promoting exon recognition, they can constrain synonymous sequence evolution [1–3]. Splicing of exitrons (exon-embedded introns) extends this principle: segments of coding sequence (CDS) are excised under the control of *cis*-regulatory information that overlaps the CDS [4]. This dual role of coding sequence is particularly relevant when the excised segment encodes an intrinsically disordered region (IDR). IDRs frequently coincide with low-complexity domains (LCDs) and mediate flexible molecular interactions, but can also promote aberrant self-association [5,6]. By removing IDR-coding sequence while leaving the amino-acid sequences of adjacent structured domains unchanged [4], exitron splicing could modulate both functional interactions and the potential for aberrant self-association. Yet IDR primary sequences often diverge rapidly, even when overall amino-acid composition or other physicochemical properties remain similar [7–11]. This divergence raises the question of whether IDR-removing exitron-type splicing recurs across species despite variation in splice-junction positions.

TDP-43, encoded by *TARDBP*, provides a tractable model for addressing this question. Its N-terminal domain (NTD) and two RNA recognition motifs (RRMs) form a structured core with comparatively conserved amino-acid sequences, whereas its sequence-divergent C-terminal IDR/LCD mediates regulatory interactions and can promote pathological self-assembly [12–17]. In humans, multiple exitron-type splicing events use donors within terminal exon 6 to remove much of the C-terminal IDR/LCD-coding sequence while leaving the NTD–RRM-coding sequence intact [18–22]. IDR-removing exitron-type splicing and splicing of introns confined to the 3′ untranslated region (3′UTR) are both TDP-43 dependent, contributing to autoregulation of full-length TDP-43 expression [23–26]. However, it remains unclear which nucleotide features support IDR-removing splicing across sequence-divergent *TARDBP* coding regions.

Here, we compared RNA-seq-supported splicing events with donors in IDR-coding sequences across human genes and their vertebrate counterparts by predicted coding outcome—IDR removal with preservation of structured regions—rather than by exact junction position. Because donor-containing sequences also encode protein, we evaluated their nucleotide features against synonymous alternatives that preserve the encoded amino acids. The teleost *TARDBP* ohnologs *tardbpa* and *tardbpb*—copies retained after whole-genome duplication [27,28]—offered a useful contrast for functional testing: compositionally similar disordered tails but different splicing patterns. In human cells, we used reciprocal donor-proximal substitutions in zebrafish minigenes to test *cis*-regulatory effects on splice-site choice and corresponding reporters to examine associated changes in protein-product balance and puncta formation. Together, these approaches distinguish the capacity to encode an IDR from the RNA-level control of its inclusion in protein products.

## Materials and Methods

### Comparative and computational analyses

#### Identification of human exitron-type splicing events predicted to remove IDRs while preserving structured regions

One complete protein-coding reference transcript per human gene was selected from Ensembl release 115 before splice-event classification, in the following priority order: MANE Select > Ensembl canonical > APPRIS principal 1–5 > longest complete translation [29–31]. Intrinsic disorder was predicted with metapredict v3.0.2 (V3 model; threshold, 0.5) [32]. Protein sequences were scanned against Pfam v38.2 [33] using HMMER v3.4 (hmmscan -- cut_ga). Annotated structured regions were defined solely by the alignment intervals of Pfam Domain and Family hits; overlapping or immediately adjacent intervals were merged. These annotations define the protected region set operationally and do not establish that every included region is stably folded in isolation. Predicted disordered segments were split at structured-region boundaries, and segments of ≥30 amino acids were declared target IDRs.

Human splice junctions were compiled from GTEx v8/recount3 via Snaptron gtexv2 (GRCh38; 17,382 RNA-seq samples from 948 participants) [34–36]. Reconstruction of post-excision sequences was attempted for all 959,375 RNA-supported gene–junction events mapped to the selected transcripts. Successfully reconstructed sequences were translated *in silico*, stopping at the first in-frame stop codon where present. An event qualified if its donor lay strictly within a single CDS segment and within a declared target IDR segment, and its reconstructed translation lacked at least 75% of the residues in the union of the declared target IDR segments within the same positional compartment (N-terminal, internal, or C-terminal) while retaining every annotated structured-region residue unchanged. Frameshifts downstream of the final annotated structured region were allowed. Acceptor annotation, termination position, predicted nonsense-mediated decay, and newly encoded C-terminal sequence were recorded but did not affect eligibility. The same exact donor–acceptor pair had to be detected in ≥100 distinct participants. Gene–junction events, gene-specific donors, and genes were counted separately. Here, exitron-type splicing refers to events with CDS-internal donors in IDR-coding regions; acceptors need not lie within the CDS or the same exon.

#### Cross-species sequence conservation and recurrence of IDR-removing splicing

Directly mapped protein-coding counterparts of the 448 human discovery-set genes were identified in mouse, chicken, coelacanth, elephant shark, spotted gar, and zebrafish using Ensembl Compara release 115 [37]. Annotated mature sequences were the default references. Alternative references retaining one coding intron required an unambiguous translation ending at the first in-frame stop codon and RNA support for both the retained form and a splice event whose donor lay within its target IDR-coding region and whose product preserved all annotated structured-region residues unchanged. Their corresponding annotated mature proteins had to match the preselected reference for that copy exactly. Among distinct retained protein sequences, the reference with the greatest predicted target-IDR removal fraction was selected; the annotated reference was retained in case of ties.

Selected proteins were aligned to their human references using MAFFT v7.526 (--globalpair --maxiterate 1000) [38], with CDS codons mapped onto the protein alignments. Identity was calculated separately for the union of donor-containing continuous IDRs and the union of annotated structured regions. Exact residue and A/C/G/T matches were divided by the full human-region lengths in residues and nucleotides, respectively; gaps and ambiguous bases contributed no matches. Both region sets required ≥80% coverage by aligned residues and by unambiguous nucleotide pairs. Identities were averaged equally across assessable copies within each human gene– species combination.

Recurrence was assessed using each copy’s own annotations and the product-reconstruction procedure described above. Qualifying reconstructed products lacked ≥75% of the donor-containing target IDR segment while preserving every annotated structured-region residue unchanged. Unlike the human analysis, removal was measured for this single segment. Each exact donor–acceptor pair required ≥1 uniquely mapped split read in each of ≥2 physical RNA-seq runs. A human gene–species combination was positive if any mapped copy contained a qualifying event; medaka was analyzed separately for sensitivity.

Local conservation was assessed across the same six species for the 45 genes with qualifying splicing in elephant shark. Human-donor-centered 50-nt coding windows (−25 to +24 nt) were projected onto target CDSs through the protein alignments while preserving codon phase. Nucleotide and amino-acid identity denominators were 50 nt and the 16 residues encoded by complete codons, respectively. Both the local window and annotated structured-region set required ≥80% coverage at both sequence levels. Within each human gene–species combination, values were averaged equally across assessable copies per donor and then across assessable donors. Paired nucleotide-identity differences were expressed in percentage points as identity in structured regions minus identity in local windows.

#### *TARDBP* comparative panels, annotations, and reference coordinates

Detailed *TARDBP* analyses included six non-teleost gnathostome loci, 15 teleost *tardbpa*/*tardbpb* pairs, and eight non-gnathostome chordate or protostome homologs. Human coordinates were based on GENCODE v39 on GRCh38 and the MANE Select transcript ENST00000240185.8 [30,39]. Gnathostome annotations were primarily from Ensembl release 115; outgroup sources included NCBI, Ensembl Metazoa, FlyBase, and WormBase [29,40,41] (Supplementary Table 1).

Assignments to *tardbpa* or *tardbpb* were prespecified using curated reference annotations and structured-core homology, then assessed for concordance with local synteny. For each teleost locus, up to ten flanking protein-coding genes on each side were retrieved from Ensembl release 115. Genes were grouped by Ensembl Compara orthology links [37], excluding *TARDBP*-family genes and genes lacking links to other sampled flanking genes. Each locus’s set of flanking-gene groups was compared separately with the pooled sets around *tardbpa* and around *tardbpb* in the other 14 species. Jaccard similarity was the number of shared groups divided by the number in the union of the two sets (Supplementary Table 2).

For each teleost copy, the coding position corresponding to the start of human exon 6 (c.715; TDP-43 residue 239) served as the coordinate anchor. The retained terminal-region reference comprised uninterrupted genomic sequence from this anchor through the first stop codon in the annotated reading frame. Candidate introns had to span ≥20 nt. CDS-internal donors lay at or downstream of the anchor and ≥10 nt upstream of the retained reference stop. For post-stop analysis only, donors and their acceptors were restricted to within 4,000 and 10,000 nt downstream of that stop, respectively. For each RNA-supported exact donor–acceptor pair, retained CDS upstream of the donor was joined to sequence downstream of the acceptor. The reconstructed sequence was translated in the annotated frame to the first stop codon to assess preservation of the NTD–RRM core sequence.

#### *TARDBP* phylogenetic and protein-sequence analyses

The human TDP-43 structured core was defined as residues 1–259, encompassing the NTD (1–76), RRM1 (106– 176), and RRM2 (191–259). For the gnathostome core phylogeny, ten proteins were aligned with MAFFT v7.526 G-INS-i, and columns containing >50% gaps were removed. A neighbor-joining tree was inferred from amino-acid p-distances in Biopython v1.87 [42], counting gap–gap positions as matches and gap–residue positions as mismatches. Branch support was assessed with 1,000 bootstrap resamples of alignment columns [43,44]. Medaka *tardbpa* and *tardbpb* were included in inference but omitted from the displayed eight-tip tree. For the tree including outgroups, regions corresponding to human residues 1–259 from 16 proteins were aligned with MAFFT --auto. The same column-filtering criterion (>50% gaps) was applied, but no columns were excluded. Pairwise distances were calculated by excluding positions containing a gap in either sequence, and neighbor-joining branch support was assessed with 200 bootstrap resamples.

For *TARDBP* and outgroup tracks, intrinsic disorder was predicted with IUPred3-long (threshold, 0.5) [45]. AlphaFold confidence was displayed as 1 − pLDDT/100 only for models whose sequences exactly matched the analyzed proteins [46–48]. Local amino-acid complexity was quantified as Shannon entropy (bits) in centered windows of up to 13 residues [49]. Positions with entropy ≤2.20 bits were provisionally classified as low complexity; this classification was retained only if their centered five-residue neighborhood contained ≥3 such positions. Proteins were aligned to human TDP-43, and CDS nucleotides were mapped onto the resulting codon axis. Amino-acid and nucleotide identity to human were summarized in 25-residue and 75-nt windows advanced by 8 residues and 24 nt, respectively.

#### Public bulk RNA-seq processing and splice-junction detection

Public short-read RNA-seq data were obtained from the Sequence Read Archive, European Nucleotide Archive, and DNA Data Bank of Japan Sequence Read Archive [50]. The primary six-species recurrence analysis used 139 physical runs, increasing to 169 with medaka in the sensitivity analysis. The teleost donor-cluster analysis used 148 runs across 15 *tardbpa*/*tardbpb* ohnolog pairs (Supplementary Table 3).

Reads from the 330 public bulk RNA-seq runs were aligned to species-specific genome assemblies using the corresponding annotations and STAR v2.7.11b (273 runs) or v2.7.3a (57 runs) [51]. Single-end or paired-end processing followed the library layout. For genome indexing, sjdbOverhang was set to 74 for the paired-end 75-nt *Ciona* larval-brain data, 149 for the paired-end 150-nt *Ciona* adult-pharynx and *Drosophila yakuba* data, and 100 otherwise. The option --outSAMstrandField intronMotif was additionally specified for the *Ciona* larval-brain and *D. yakuba* data. Other splice-junction filtering, alignment-overhang, and multimapping parameters were left at their defaults for the STAR version used.

Exact donor–acceptor pairs were identified from SJ.out.tab using strand information and recorded separately for each run. A junction was considered detected in a run when supported by ≥1 uniquely mapped split read; multimapping reads were excluded from support counts. For quantification, junction-read counts were obtained from STAR junction files, whereas reads spanning the donor boundary within one contiguous alignment block were counted from indexed BAM alignments.

#### Donor-cluster spliced-read fraction and retained-interval coverage

Eligible junctions were grouped by donor. For each locus and donor class, the cluster with the greatest pooled uniquely mapped STAR junction-read support across the fixed run set was selected before calculating fractions or sequence scores. Within selected CDS-internal clusters, acceptors were ranked by pooled junction-read support.

The donor-cluster spliced-read fraction was calculated per run as *J*/(*J* + *S*), where *J* was the number of uniquely mapped STAR junction reads sharing the donor and *S* was the number of distinct query names of reads spanning the donor boundary within one contiguous alignment block. *S* was calculated using only primary, uniquely mapped, nonduplicate, nonsupplementary alignments. Locus-level fractions were arithmetic means of run-level fractions; CDS-internal comparisons required an eligible cluster at both loci.

For *tardbpb* loci lacking an eligible CDS-internal cluster, coverage was assessed from the coding anchor through the retained reference stop codon. Per-base depth was calculated by counting distinct query names for reads covering each base within aligned blocks, excluding spliced gaps and using the same alignment filters as for *S*. Complete-interval coverage required depth ≥1 at every base in every analyzed run. Fractions, pooled CDS-internal junction-read counts, and coverage summaries are provided in Supplementary Table 4.

#### Outgroup terminal-region survey

For *Drosophila* and *Caenorhabditis* homologs, each annotated intron in the C-terminal coding region was individually reinserted into the mature CDS *in silico*. The reconstructed sequence was translated in the annotated reading frame to the first stop codon to assess the coding consequence of retention. Junctions from terminal CDS-internal donors in *Ciona*, *Branchiostoma*, and *Petromyzon* were surveyed using STAR alignments. Reproducibility required ≥1 uniquely mapped split read supporting the same exact donor–acceptor pair in each run of a same-tissue replicate pair.

#### Splice-site strength of human IDR-removing events

Canonical GT donors and AG acceptors were scored using MaxEntScan score5 (−3 to +5 nt from donor G) and score3 (−19 to +3 nt from acceptor G), respectively [52]. Controls were within-gene conventional introns locally constitutive across complete protein-coding transcripts spanning the site, with donors absent from the human CDS-internal donor set and coding boundaries ≥10 amino acids inside annotated structured regions (1,276 controls). For donors, pre-stop acceptors, and post-stop acceptors, effects were mean gene-level differences between focal and control median scores (focal minus control). Percentile 95% confidence intervals and two-sided *P* values were calculated using 10,000 gene bootstrap resamples and 10,000 gene-level sign-flip permutations, respectively; *P* values were Holm-corrected across the three contrasts.

Donor comparisons were repeated by target-IDR position relative to annotated structured regions (N-terminal, internal, or C-terminal), using the same estimator and resampling procedures with Holm correction across strata. Each eligible gene contributed to only one stratum. Between-stratum heterogeneity was assessed by permuting stratum labels among genes while preserving stratum sizes, using the maximum absolute studentized pairwise mean difference. Pairwise comparisons used 10,000 label permutations for *P* values and 10,000 independent gene bootstrap resamples for confidence intervals, with Holm correction across the three comparisons.

#### Synonymous-sequence compatibility of human CDS-internal donors

Each donor position was analyzed once per gene, irrespective of the acceptor. Canonical GT donors were eligible if the entire 9-nt score5 window lay within a single retained coding exon and all overlapping codons encoded residues in the same continuous target IDR. These criteria retained 458 donors from 381 genes.

Controls were other GT sites in the same target IDR that met the same coding, exon-continuity, and codon-completeness criteria. Candidate controls were excluded if they coincided with any discovery-set donor, were annotated as donors in Ensembl release 115, or had a score5 window overlapping a discovery-set donor dinucleotide. This yielded 14,083 control GTs; one gene lacked an eligible control, leaving 380 genes for paired analysis.

For each focal and control site, all distinct 9-mers obtainable by synonymous recoding of the complete codons overlapping the score5 window were enumerated, preserving the local peptide and GT at positions 0 and +1. Each resulting 9-mer was scored with MaxEntScan score5. The native sequence’s midrank percentile was

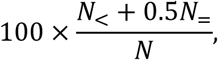

where *N* is the number of distinct scored 9-mers, and *N*_<_ and *N*_=_ count those scoring below and equal to the native 9-mer, respectively. Within each gene, focal and control percentiles were summarized separately by their medians. The primary estimate was the median across genes of the paired difference between these medians (focal minus control). Its 95% confidence interval was calculated using 10,000 gene bootstrap resamples. An exact two-sided sign test evaluated the paired differences, excluding zeros. A sensitivity analysis restricted controls to coding phases represented among eligible focal donors in the same IDR.

#### Donor-centered nucleotide conservation across vertebrates

Non-human GT and GC donors were included when at least one exact junction met the coding-outcome criteria and was detected in ≥2 physical RNA-seq runs (131 donor observations; 47 human genes). Each window comprised 16 complete codons, with phase-dependent boundaries collectively spanning −26 to +23 nt relative to donor G (position 0). At directly aligned coding bases, identity to human was compared with an analytic synonymous expectation conditioned on the target copy’s amino-acid sequence and all active donor dinucleotides. Synonymous codons were weighted by copy-specific CDS usage (pseudocount, 1 per codon). All donor-dinucleotide positions constrained in the model were excluded from identity summaries.

For each summary, aligned-base observations were averaged within human gene–species combinations, then equally across species within genes and across genes. Identity excess (observed minus expected identity; percentage points) was summarized by donor-relative position, four flanking intervals (−26 to −13, −12 to −1, +2 to +12, and +13 to +23 nt), and codon position. The 95% confidence intervals were calculated using 10,000 gene bootstrap resamples; two-sided *P* values were calculated using 10,000 sign-flip permutations of complete gene profiles. Benjamini–Hochberg (BH) correction [53] was applied separately to the three test families. Analyses were repeated excluding *TARDBP*.

#### *TARDBP* splice-site scores and donor-and acceptor-flanking sequence profiles

Six loci were analyzed: human, mouse, coelacanth, spotted gar, elephant shark, and zebrafish *tardbpa*. MaxEntScan score5 was calculated for 27 conventional-intron and 13 CDS-internal GT donors, and score3 for 26 conventional-intron AG acceptors and 14 AG acceptors linked to CDS-internal donors. One shared acceptor was excluded from the conventional-intron set. Each comparison used an exact two-sided sign test on six pairs of locus medians.

The donor profile used 13 CDS-internal GT-centered 50-nt windows (−25 to +24 relative to donor G); the 12 non-teleost sites formed the Fig. 4 reference. Frequencies with a pseudocount of 0.5 per base were converted to log₂ odds against a uniform background. Window scores were calculated by summing these values over A/C/G/T positions. Controls were 84 GT sites matched by species and coding phase in the corresponding terminal CDSs, excluding observed donors.

An exploratory 1–4-nt k-mer screen used 16 donors, 107 matched GT controls, and 100,000 species/locus-stratified GT-background resamples, with BH correction [53] across 1,414 k-mer–position pairs. Ten significant pairs (q ≤ 0.05) within −25 to +24 were manually selected to represent five sequence blocks, considering GT-background and codon-aware support. Final comparisons used an overlapping cohort of 13 donors and 84 controls, together with 10,000 synonymous profiles preserving amino-acid sequences and sampling from all synonymous codons according to reference-specific terminal-CDS usage. One-sided Fisher exact tests compared observed k-mer counts with GT-control or pooled synonymous-background counts, with separate BH correction across the ten pairs for each comparison.

The acceptor profile used 14 AG sites directly linked to CDS-internal donors, with complete windows spanning −50 to +50 nt relative to acceptor G; the 13 non-teleost sites formed the Fig. 4 reference. Distal sites attributable to serial downstream processing were excluded. Each site was matched to the 20 nearest post-stop AG controls whose complete windows did not overlap any compiled acceptor window. The a priori alternative of a greater mean C/T fraction at observed sites (−50 to −3) was tested using 100,000 site-stratified permutations, each sampling one control per observed site; *P* values were calculated with add-one correction. Species-balanced and leave-one-species-out analyses assessed sensitivity.

#### Teleost donor, acceptor, and exact-coordinate comparisons

Donor clusters were selected as described above. For each selected CDS-internal cluster, the observed acceptor was the site with the greatest summed unique junction-read support across the fixed run set. Observed *tardbpa* donor windows were paired with codon-projected *tardbpb* windows. Uninterrupted anchor-to-stop sequences were translated, and the resulting proteins were globally aligned (match, +2; mismatch, −1; gap, −2). The *tardbpa* donor codon and within-codon offset were projected onto *tardbpb*; when the codon aligned to a gap, the nearest aligned *tardbpb* codon was used and flagged. Paired 50-nt windows (−25 to +24) were scored against the 12-site non-teleost donor profile. Window scores were also compared between observed GT donors in the ten pairs with eligible CDS-internal clusters at both loci.

Acceptor matching used transcript-oriented 61-nt windows (−50 to +10 relative to acceptor G). Within ±600 nt of the corresponding stop-relative position in *tardbpb*, the canonical AG-centered window with the highest nucleotide identity to the observed *tardbpa* acceptor window was selected; the maximum was unique in all 15 pairs. These sequence-matched candidate regions were not assumed to be orthologous, native, or RNA-supported acceptors. Paired windows were scored using positions −50 to +10 of the 13-site non-teleost acceptor profile.

Exact-human-coordinate controls were analyzed separately in each ohnolog. Human c.842 was projected through MAFFT protein alignments (--globalpair --maxiterate 1000), preserving codon correspondence and within-codon offset. The G of the major human acceptor, 538 nt downstream of the reference stop, was projected through pairwise MAFFT alignments of 2-kb terminal-anchor nucleotide sequences using the same options. Contiguous 50-nt windows (−25 to +24) were extracted around the projected coordinates without local realignment. Identity to the corresponding human windows was calculated only over positions containing A/C/G/T in both sequences.

MaxEntScan score5 at each RNA-seq-selected *tardbpa* donor was compared with the highest score among canonical GT candidates within −25 to +24 nt of its nucleotide-projected position in *tardbpb*. The search was repeated over −40 to +40 nt for sensitivity analysis. Pairs without a canonical GT in the relevant *tardbpb* interval were excluded from that comparison.

#### Teleost post-RRM2 protein composition and translation-conditioned donor-profile scores

For each copy in the 15 *tardbpa*/*tardbpb* pairs, the RRM2 endpoint was the last residue aligned to human TDP-43 residues 192–259 in a global BLOSUM62 alignment (gap opening, −10; gap extension and terminal gaps, −0.5). The post-RRM2 region comprised all subsequent residues. Composition similarity was 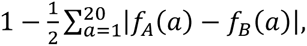 where *f_A_* and *f_B_* are normalized frequencies of the 20 standard amino acids in the paired post-RRM2 regions. Residue identity was the fraction of exact matches among alignment columns containing residues from both copies.

In each copy, a 50-nt window (−25 to +24) was centered 85 nt downstream of the coding anchor defined above. This position coincided with an RNA-supported donor in every *tardbpa* copy. For each copy, 50,000 null sequences were generated by permuting codons without replacement among positions encoding the same amino acid, preserving the amino-acid sequence and counts of each codon. Native and null sequences were scored against the 12-site non-teleost donor profile, excluding positions 0 and +1 from scoring. Donor-profile score excess was the native score minus the median null score. Prespecified sensitivity analyses varied profile masking, focal-codon treatment, and reference-site weighting.

#### RBPmap prediction of TDP-43 motif matches

Transcript-oriented sequences from +1 to +4,000 nt downstream of the reference stop codon were submitted to RBPmap using the “other” genome setting, default stringency, and *TARDBP* motif models 284 and 285 [54]. Paired *tardbpa*/*tardbpb* comparisons used predicted matches within the first 2,000 nt. The primary summary was the sum of server-reported Z scores per kilobase of analyzed sequence. Secondary summaries were predicted-match density per kilobase, the sum of the three highest Z scores, and the sum of −log₁₀ *P* values per kilobase.

#### Zebrafish single-cell RNA-seq expression analysis

Expression of zebrafish *tardbpa* and *tardbpb* was analyzed in ten processed adult-brain loom matrices from GSE246714 [55] (Supplementary Table 3), using the dataset labels *tardbpl* and *tardbp*, respectively. Detection was defined as a raw unique molecular identifier (UMI) count >0. Detection fractions were calculated for each gene separately, both genes in the same cell, and at least one of the two genes, without normalization or pseudocounts.

For expression summaries, counts were normalized per cell to 10,000 total UMIs (CP10K). Within each GEO sample, cells were grouped by the supplied cluster labels; groups with ≥20 cells were retained. For each gene, mean CP10K was calculated within each group and then transformed as log₂(mean CP10K + 0.01). Pearson and Spearman correlations between the two genes were calculated across the 108 retained sample–cluster groups.

### Functional assays

#### Minigene and reporter construction

The human *TARDBP* minigene spanning exons 5–6 was cloned into pcDNA3.1/myc-His(−)A. Elephant shark, spotted gar, and zebrafish *tardbpa*/*tardbpb* constructs were designed from Ensembl transcript models. Transcript-oriented inserts began within the coding region corresponding to human *TARDBP* exon 5 and included the intervening intron, terminal coding exon, 3′UTR, and downstream regulatory sequence. The endpoint was generally the more distal of the annotated transcript end and the position 100 bp beyond the last retained RBPmap-predicted motif cluster, adjusted to avoid neighboring genes or plausible RNA-processing signals. Constructs were synthesized by VectorBuilder and expressed from a cytomegalovirus (CMV) promoter.

Reciprocal zebrafish substitutions replaced the donor dinucleotide, selected flanking sequence, or the −25 to +24 or −40 to +40 interval with aligned sequence from the other ohnolog, leaving the remaining sequence unchanged. gBlocks fragments (Integrated DNA Technologies) and inverse-PCR backbones were assembled with In-Fusion Snap Assembly. Reporters carried in-frame EGFP immediately after the initiation codon. The synonymous AGT→TCT donor-disrupting *tardbpa* reporter was generated using the KOD-Plus-Mutagenesis Kit, DpnI digestion, and self-ligation. Construct coordinates, identifiers, sequences, exchanged intervals, synthetic fragments, primers, and siRNAs are provided in Supplementary Table 5.

#### Cell culture, transfection, and TDP-43 perturbation

HEK293T cells (KAC, EC12022001-F0) were maintained in Dulbecco’s modified Eagle’s medium (Gibco, 11995065) with 10% fetal bovine serum (Gibco, 10270106) at 37 °C in 5% CO₂. For TDP-43 perturbation (Fig. 2), cells were seeded at 5 × 10⁴ per well in 24-well plates 24 h before cotransfection with 100 ng minigene, 50 ng EGFP–TDP-43 (VectorBuilder, VB240423-1225beg) or EGFP control plasmid, and 10 nM endogenous *TARDBP* 5′UTR-targeting or non-targeting siRNA. Lipofectamine 3000 (Invitrogen, L3000008) was used in Opti-MEM without P3000. The siRNA target was absent from the EGFP–TDP-43 construct, and total plasmid DNA was 150 ng per well. Reciprocal-substitution minigenes (Fig. 5) were assayed under basal TDP-43 conditions using 100 ng minigene per well. Cells in both assays were harvested 48 h after transfection.

#### RNA extraction and splice-product quantification

Total RNA was isolated with the NucleoSpin RNA kit (Macherey-Nagel, 740955.10), including on-column rDNase treatment, and quantified with a NanoDrop One. Reverse transcription used 500 ng RNA in 10 µL with PrimeScript RT Master Mix (Perfect Real Time; TaKaRa, RR036A) at 37 °C for 15 min; cDNA was diluted to an RNA-equivalent concentration of 2 ng/µL.

Endpoint RT-PCR used TaKaRa LA Taq (RR002A) in 25-µL reactions containing 10 ng RNA-equivalent cDNA. Cycling comprised 94 °C for 1 min, followed by 31 cycles of 94 °C for 30 s, 60 °C for 30 s, and 72 °C for 1 min, with a final extension at 72 °C for 5 min. Vector-specific primers distinguished human minigene transcripts from endogenous *TARDBP*; construct-specific primers were used for non-human minigenes (Supplementary Table 5). Products were resolved on 2% agarose gels containing GelRed (Biotium, 41003), imaged on an Amersham Imager 680 (Cytiva), and quantified with ImageQuant. *In* and *Ex* denote IDR-retaining and IDR-removing products, respectively.

RT-qPCR used TB Green Premix Ex Taq II (TaKaRa, RR820R) on a Thermal Cycler Dice Real Time System III (TaKaRa Bio, TP950), in technical duplicate, using 25-µL reactions containing 10 ng RNA-equivalent cDNA. Cycling comprised 95 °C for 30 s and 40 cycles of 95 °C for 5 s and 60 °C for 30 s, followed by dissociation analysis at 95 °C for 15 s, 60 °C for 30 s, and 95 °C for 15 s. Junction-spanning assays avoided unspliced precursor amplification. Specificity was assessed by melt-curve analysis, and only primer pairs with validated amplification efficiencies were used.

For Fig. 2, *Ex*/*In* was determined by endpoint RT-PCR for the human minigene and RT-qPCR for non-human minigenes. For RT-qPCR, *In* and *Ex* were normalized to the same minigene-RNA reference assay, and relative abundances were calculated using the 2^−ΔΔCt^ method relative to the matched control before calculating *Ex*/*In*. Ratios from both methods were expressed as log₂ fold change relative to control. For Fig. 5, *In* was normalized to the constitutive-junction (*CJ*) signal. This assay detects transcripts containing the upstream constitutive exon–exon junction and excludes *bypass* transcripts. *In*/*CJ* was expressed as log₂ fold change relative to the mean *In*/*CJ* of the corresponding unsubstituted control group. Endpoint RT-PCR ratios were expressed as log₂(*Ex*/*In*) after summing the indicated *Ex* bands and excluding *bypass*. Unsubstituted controls were excluded from *Ex*/*In* quantification because *In* (*tardbpa*) or *Ex* (*tardbpb*) was undetectable by endpoint RT-PCR.

### Immunoblotting and assignment of reporter-product bands

For reporter immunoblotting, HEK293T cells were seeded at 1 × 10⁵ per well in 24-well plates, cotransfected with 100 ng EGFP reporter and 50 ng mCherry plasmid (VectorBuilder, VB010000-9835xck), and harvested after 48 h. For immunoblotting, cells were lysed in RIPA buffer (Nacalai Tesque, 16488-34) with protease inhibitors (Sigma-Aldrich, P8340), sonicated, and centrifuged at 20,000 × g for 30 min at 4 °C. RIPA-soluble protein was quantified by BCA assay (FUJIFILM Wako, 297-73101). Samples (7.5 µg per lane) were separated by 10% SDS–PAGE and transferred to PVDF at 100 V for 1 h at 4 °C using wet tank transfer. Total protein was visualized with No-Stain reagent (Thermo Fisher Scientific, A44449).

Membranes were blocked in 5% skim milk for 1 h at room temperature and incubated with primary antibodies overnight at 4 °C: anti-TDP-43 (Proteintech, 12892-1-AP; 1:5,000) for Fig. 2, anti-GFP (Proteintech, 50430-2-AP; 1:2,000), or anti-mCherry (Abcam, ab167453; 1:1,000). Detection used HRP-conjugated anti-rabbit IgG (Dako, P0448). mCherry was analyzed on separate membranes using aliquots of the same lysates. Signals were developed with Immobilon Western substrate (Millipore, WBKLS0500), acquired on an Amersham Imager 680, and quantified with ImageQuant.

TDP-43 signals were normalized to total protein. GFP and mCherry signals were normalized to total protein on their respective membranes and to a common reference sample included on every gel. Relative reporter-protein abundance in the RIPA-soluble fraction was expressed as normalized GFP-band intensity relative to the normalized mCherry signal. GFP-reactive bands were assigned operationally from corresponding RNA products, apparent molecular masses, and responses to *In*-selective siRNAs (Supplementary Table 6). These siRNAs targeted sequence retained in *In* but removed by IDR-removing splicing; siRNA-sensitive bands were classified as *In*-associated.

### Live-cell imaging and selective depletion of the IDR-retaining transcript

HEK293T cells were seeded in µ-Slide 8 Well chambers (ibidi, ib80826) and transfected the next day with 100 ng EGFP reporter and 30 ng mCherry plasmid using 0.3 µL Lipofectamine 3000 in Opti-MEM without P3000. *In*-selective depletion assays used 100 ng reporter, 50 ng mCherry plasmid, and 20 nM cognate or non-targeting siRNA. At 48 h, cells were stained with Hoechst 33342 (Invitrogen, H3570; 1:10,000) for 10 min at 37 °C and imaged live on a Keyence BZ-X810 with a 20× objective.

Whole-cell segmentation used Fiji/ImageJ (2.18.0/1.54p) [56], the Fiji-Cellpose plugin (Cellpose_Appose v0.1.17; imglib2-cellpose v0.0.8), and the Cellpose 3 cyto3 model [57,58], using mCherry and Hoechst but not EGFP. mCherry positivity was determined from mean raw whole-cell intensity, including the nucleus, without background subtraction or normalization. A single absolute threshold established from representative images before analysis was applied across all conditions and replicates. Puncta were scored manually. A puncta-positive cell was an mCherry-positive cell containing at least one discrete cytoplasmic EGFP focus. Foci at cell boundaries were reviewed and assigned to one cell only. Dead or morphologically abnormal cells, cells with unassignable overlapping puncta, and cells with saturated nuclear EGFP were not classified as puncta-positive.

Counts were recorded separately for each of four randomly selected fields per well. For well-level summaries, counts were pooled across fields, and the puncta-positive fraction was calculated as the number of puncta-positive cells divided by the total number of mCherry-positive cells. Image acquisition was unblinded; puncta scoring was blinded to construct identity.

### Statistical analyses

Within-species ohnolog pairs were the units for teleost comparisons. Three prespecified Fig. 4 tests evaluated *tardbpa* > *tardbpb*: eligible CDS-internal donor-cluster detection (15 pairs; exact binomial test on discordant pairs), donor-cluster spliced-read fractions (10 pairs), and donor-profile scores for observed *tardbpa* versus codon-projected *tardbpb* windows (15 pairs). The latter two comparisons used exact paired sign tests; all three tests were one-sided, and their *P* values were Holm-adjusted as one family (α = 0.05) [59]. Descriptive secondary paired comparisons used unadjusted exact two-sided sign tests; translation-conditioned donor-profile excess was evaluated using a separate unadjusted exact one-sided paired sign test under the same alternative. Ties were excluded from sign tests. These tests summarize directional consistency among sampled pairs and do not imply that the pairs represent phylogenetically independent evolutionary events. Analysis-specific resampling and false-discovery-rate procedures are described in the corresponding sections.

Cell-based assays used culture wells as biological replicates (*n* = 4), with imaging fields and cells as subsamples. RT-qPCR and endpoint RT-PCR measurements were analyzed using one-way ANOVA with Tukey’s tests for all pairwise comparisons. Immunoblot comparisons with the indicated reference used two-sided t-tests based on the common residual variance from one-way ANOVA, with Holm correction within each reporter background and protein product. Skewed or heteroscedastic ratios and fold changes were log₂-transformed before ANOVA; undetected bands were treated as missing. Field-level puncta-positive cell counts and mCherry-positive totals were analyzed using beta-binomial mixed models with a logit link and a random intercept for well (glmmTMB [60]; R v4.2.0). Comparisons with the corresponding parental reporter used Dunnett-type contrasts with single-step multivariate-t adjustment. Ordered substitution trends used one-sided Jonckheere–Terpstra tests, excluding the synonymous TCT construct, with Holm correction across the two reporter backgrounds. Two-group comparisons used two-sided Welch’s *t*-tests. Unless otherwise specified, tests were two-sided with *P* < 0.05 considered significant, using adjusted *P* values where applicable.

## Results

### IDR-removing exitron-type splicing recurs despite sequence divergence in donor-containing IDRs

To identify human exitron-type splicing events predicted to remove IDRs, we examined junctions with CDS-internal donors in IDR-coding regions. We reconstructed the coding consequences of exact junctions detected in at least 100 of 948 GTEx participants and retained events predicted to remove ≥75% of the target IDR residues while leaving every annotated structured-region residue unchanged. These criteria identified 820 gene–junction events involving 545 gene-specific donors in 448 genes (Fig. 1A).

**Figure 1.**
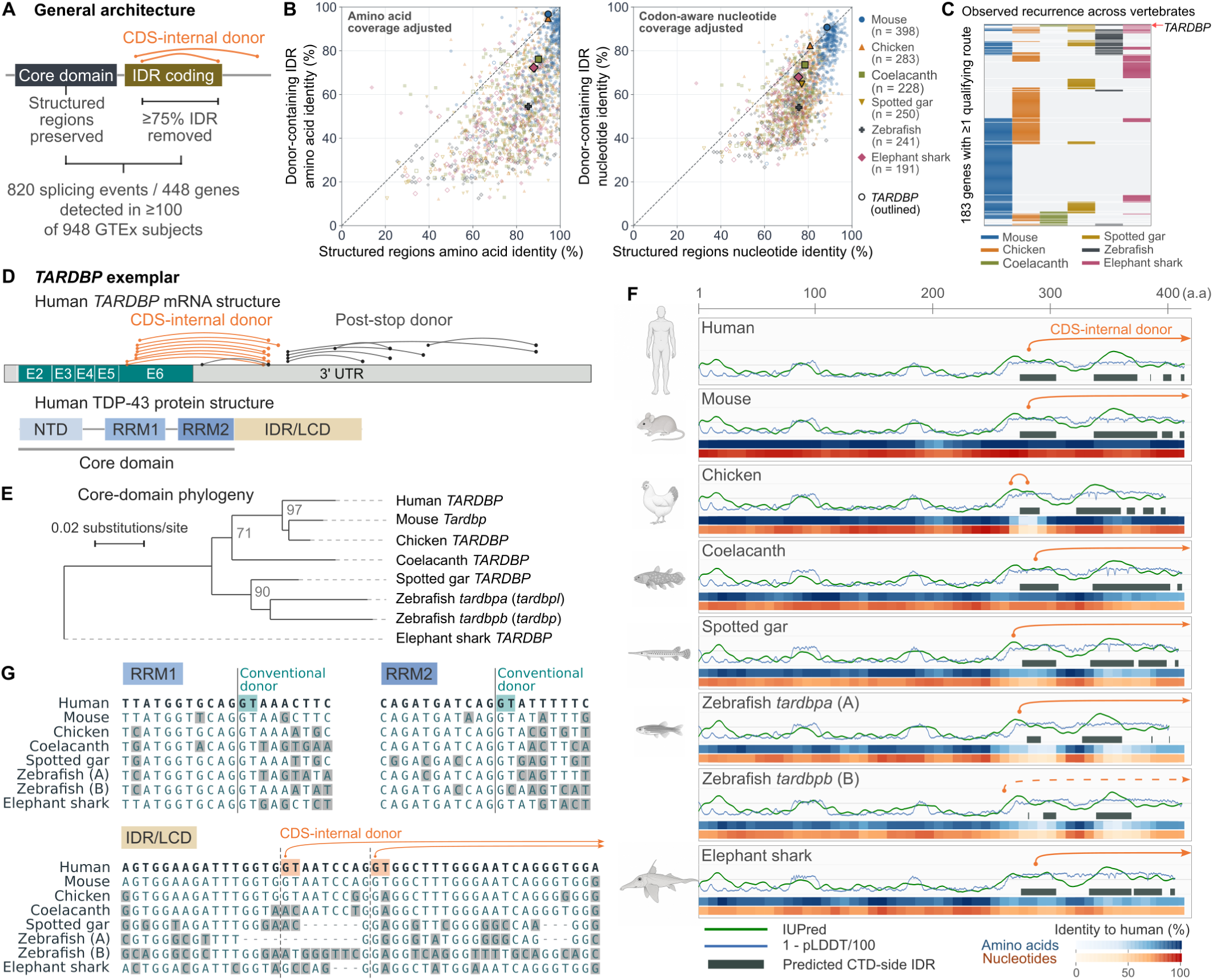
Exitron-type splicing predicted to remove IDRs while preserving structured regions recurs across jawed vertebrates. **(A)** Human CDS-internal splicing events detected in ≥100 of 948 GTEx participants, with reconstructed translations lacking ≥75% of the target IDR residues while retaining all annotated structured-region residues unchanged (820 gene–junction events; 545 gene-specific donors; 448 genes). **(B)** Coverage-adjusted amino-acid (left) and codon-aware nucleotide (right) identity to human in annotated structured regions versus donor-containing IDRs across six non-human species. Points represent assessable human gene–species comparisons; colors denote species, black outlines mark *TARDBP*, and dashed lines indicate equal identity. **(C)** Recurrence across 183 genes with qualifying splicing in ≥1 non-human species. Colored cells indicate ≥1 qualifying exact junction detected in ≥2 RNA-seq runs. Blank cells indicate unassessable combinations or combinations without qualifying RNA-seq support. **(D)** Human *TARDBP* transcript and TDP-43 domain organization. Coding regions are teal and the 3′UTR gray; orange and black arcs indicate junctions from CDS-internal and post-stop donors, respectively. **(E)** Neighbor-joining tree of jawed-vertebrate TDP-43 structured cores corresponding to human residues 1–259. Bootstrap support ≥70% is shown; scale bar, amino-acid p-distance. **(F)** Human-aligned tracks showing IUPred3 disorder scores, inverted AlphaFold confidence (1 − pLDDT/100), predicted C-terminal IDRs, and amino-acid and nucleotide identity to human. Orange arcs indicate CDS-internal splicing; the dashed zebrafish *tardbpb* arc denotes low read support, and arrowheads indicate junctions to post-stop acceptors. **(G)** Protein-guided codon alignments around conventional RRM1/RRM2 intron boundaries and the region corresponding to human exon 6 donors c.833 and c.842. Figure 1 alt text: Schematics, sequence comparisons, and a heat map summarize predicted removal of intrinsically disordered regions across surveyed vertebrate species and illustrate variable splice positions in TARDBP.

Across six non-human vertebrate species, we compared amino-acid and nucleotide identity to human between donor-containing IDRs and annotated structured regions within the same genes. Among 1,591 sequence-assessable gene–species comparisons from 416 genes, structured regions showed higher identity than donor-containing IDRs in 1,488 amino-acid comparisons (93.5%) and 1,451 codon-aware nucleotide comparisons (91.2%; Fig. 1B).

To assess cross-species recurrence of the predicted coding outcome, we searched other vertebrates for splicing predicted to remove ≥75% of the donor-containing target IDR segment while leaving every annotated structured-region residue unchanged, without requiring conservation of junction coordinates, splice phase, or termination site. Of the 448 human discovery-set genes, 435 had directly mapped protein-coding counterparts in at least one surveyed species, and 334 had at least one counterpart assessable for splicing. Qualifying junctions supported in at least two RNA-seq runs were detected in counterparts of 183 genes in at least one species, 82 genes in at least two species, and 36 genes in at least three species (Fig. 1C). Because RNA-seq coverage and sampling depth varied across species, these counts reflect observed recurrence.

At a finer scale, we examined sequence conservation in 50-nt coding windows centered on human donor positions, focusing on the 45 genes with qualifying splicing in elephant shark, the earliest-diverging lineage in our six-species panel. In the human–elephant shark comparison, 30 of the 39 sequence-assessable genes showed lower nucleotide identity in these windows than in annotated structured regions (Supplementary Fig. 1). Together, these findings indicate that predicted IDR removal recurs across vertebrates despite sequence divergence in both donor-containing IDRs and regions aligned to human donor neighborhoods.

### A core-preserving splice architecture recurs at different junctions in jawed-vertebrate *TARDBP*

To examine this recurrence in relation to transcript and protein architecture, we focused on *TARDBP*. The structured TDP-43 core, comprising the NTD, RRM1, and RRM2, showed high amino-acid sequence identity to human across the jawed-vertebrate panel (Fig. 1D–F). By contrast, the C-terminal IDR/LCD and its coding sequence showed markedly lower amino-acid and codon-aware nucleotide identity to human, respectively (Fig. 1F). This regional contrast was also evident in local alignments: coding sequences flanking the conventional RRM1 and RRM2 introns remained alignable, whereas the region corresponding to the human c.833 and c.842 donor sites contained extensive nucleotide differences and alignment gaps (Fig. 1G).

Despite this divergence, RNA-seq supported CDS-internal splicing at all eight jawed-vertebrate loci displayed in Fig. 1F, although read support was low for zebrafish *tardbpb*. At seven of these loci, at least one CDS-internal donor was joined to an acceptor downstream of the reference stop codon. Reconstructions at these loci predicted loss or replacement of much of the native C-terminal region. The chicken locus instead showed a 48-nt excision within the C-terminal IDR-coding region. Across the seven loci with post-stop acceptors, this predicted coding outcome recurred despite variation in donor position, splice phase, acceptor position, and termination site.

We surveyed eight non-gnathostome homologs to place this architecture in a broader phylogenetic context. The annotated introns in the C-terminal coding regions of four drosophilid *TBPH* loci and *Caenorhabditis elegans tdp-1* introduced premature stop codons when retained *in silico*, consistent with conventional coding-region introns (Supplementary Fig. 2A–C). In *Ciona*, *Branchiostoma*, and *Petromyzon*, neither annotations nor the surveyed RNA-seq data revealed reproducible splicing from coding sequence downstream of the position corresponding to the start of human *TARDBP* exon 6. All eight homologs nevertheless contained IUPred3-predicted disorder-enriched post-RRM2 regions (Supplementary Fig. 2D). Elephant shark was the earliest-diverging positive comparator in the sampled panel.

### Sequence-divergent *TARDBP* terminal regions support TDP-43-responsive IDR-removing splicing

We next asked whether sequence-divergent jawed-vertebrate *TARDBP* terminal regions could support TDP-43-responsive IDR-removing splicing in a common human *trans* environment. We examined human *TARDBP*, its spotted gar and elephant shark orthologs, and zebrafish *tardbpa*. The non-human post-stop regions showed limited nucleotide identity to human, yet all four loci contained RBPmap-predicted matches to UG-rich TDP-43 motifs (Fig. 2A). For functional testing, we constructed minigenes spanning the human *TARDBP* exon 5-equivalent coding region, intervening intron, terminal coding exon, 3′UTR, and downstream regulatory sequence.

**Figure 2.**
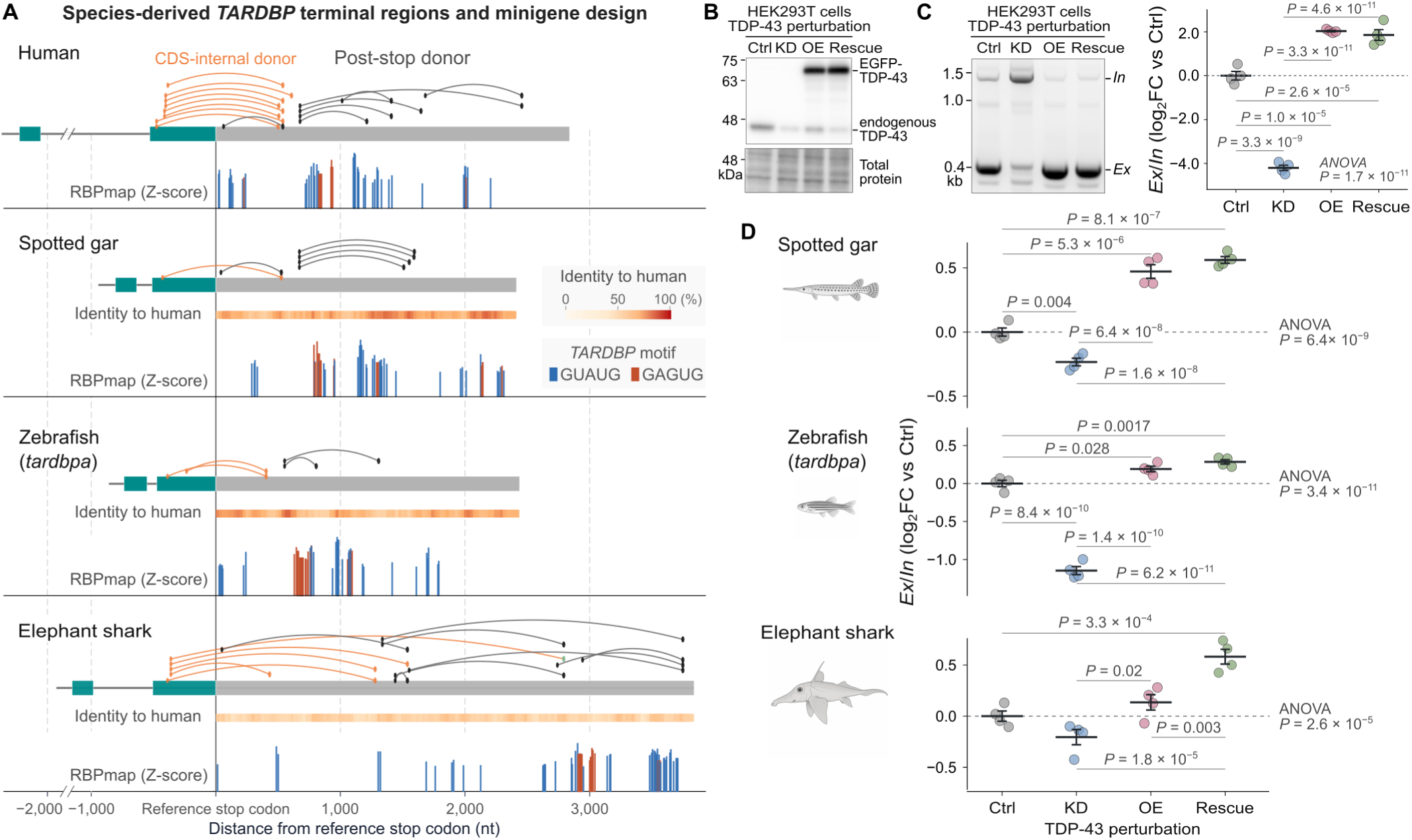
Sequence-divergent *TARDBP* terminal regions support TDP-43-responsive IDR-removing splicing. **(A)** Minigene cassettes aligned at their reference stop codons, spanning the human *TARDBP* exon 5-equivalent coding region, intervening intron, terminal coding exon, 3′UTR, and downstream regulatory sequence. Orange and black arcs indicate junctions from CDS-internal and post-stop donors, respectively. Heat maps show nucleotide identity to human; vertical bars mark RBPmap-predicted TDP-43 motif matches, with height indicating Z score. **(B)** Representative immunoblot of HEK293T cells under the following conditions: control (Ctrl), endogenous TDP-43 knockdown (KD), EGFP–TDP-43 overexpression (OE), and KD plus siRNA-resistant EGFP–TDP-43 rescue. Total-protein staining serves as a loading control. **(C)** Vector-specific endpoint RT-PCR of human *TARDBP* minigene transcripts, resolving IDR-retaining inclusion (*In*) and IDR-removing exclusion (*Ex*) products. *Ex*/*In* was quantified from band intensities. **(D)** Construct-specific RT-qPCR of spotted gar, zebrafish *tardbpa*, and elephant shark minigenes under the same conditions. In **C** and **D**, *Ex*/*In* ratios are expressed as log₂ fold change relative to control; bars show mean ± SEM (*n* = 4). Omnibus *P* values are from one-way ANOVA; pairwise *P* values are Tukey-adjusted. Figure 2 alt text: Minigene diagrams, gels, and quantitative plots show lower mean exclusion-to-inclusion ratios after TDP-43 knockdown and higher mean ratios after overexpression across four vertebrate constructs in HEK293T cells.

Each minigene was transfected into HEK293T cells under four conditions: control, endogenous TDP-43 knockdown, EGFP–TDP-43 overexpression, and knockdown combined with siRNA-resistant EGFP–TDP-43 rescue. Immunoblotting confirmed depletion of endogenous TDP-43 after knockdown and expression of EGFP– TDP-43 in the overexpression and rescue conditions (Fig. 2B). EGFP–TDP-43 overexpression was also accompanied by a lower endogenous TDP-43 signal, consistent with autoregulatory suppression.

Because RT-qPCR could not distinguish human minigene-derived transcripts from endogenous *TARDBP* transcripts, we used vector-specific endpoint RT-PCR to resolve IDR-retaining inclusion (*In*) and IDR-removing exclusion (*Ex*) products. The *Ex*/*In* band-intensity ratio was expressed as log₂ fold change relative to control. Mean values were lower under TDP-43 knockdown than under control conditions and higher under overexpression or rescue than under knockdown alone (Fig. 2C). Construct-specific RT-qPCR showed the same directional changes in *Ex*/*In* for the spotted gar, zebrafish *tardbpa*, and elephant shark minigenes, although response magnitudes varied among constructs (Fig. 2D). Thus, sequence-divergent *TARDBP* terminal regions supported TDP-43-responsive changes in the balance of IDR-retaining and IDR-removing transcripts in human cells.

### IDR-removing donors have low predicted splice-site strength but rank highly among synonymous alternatives

We next investigated the sequence basis of IDR-removing splicing, beginning with local splice-site scores in the human discovery set. IDR-removing GT donors had lower MaxEntScan score5 values than conventional intron donors whose coding boundaries lay within annotated structured regions of the same genes (275 genes; mean difference between within-gene medians, −5.663; 95% bootstrap CI, −6.409 to −4.954; Holm-adjusted *P* = 3.0 × 10⁻⁴; Fig. 3A). The magnitude of this difference varied with target-IDR position relative to the annotated structured regions (before the first structured region [N-terminal], between structured regions [internal], or after the last [C-terminal]; omnibus *P* = 0.029), with the largest reduction for C-terminal IDRs. AG acceptors before and after the reference stop codon likewise had lower score3 values than their corresponding conventional controls (237 and 74 genes; mean differences, −2.550 and −2.911; Holm-adjusted *P* = 3.0 × 10⁻⁴ and 4.0 × 10⁻⁴, respectively).

**Figure 3.**
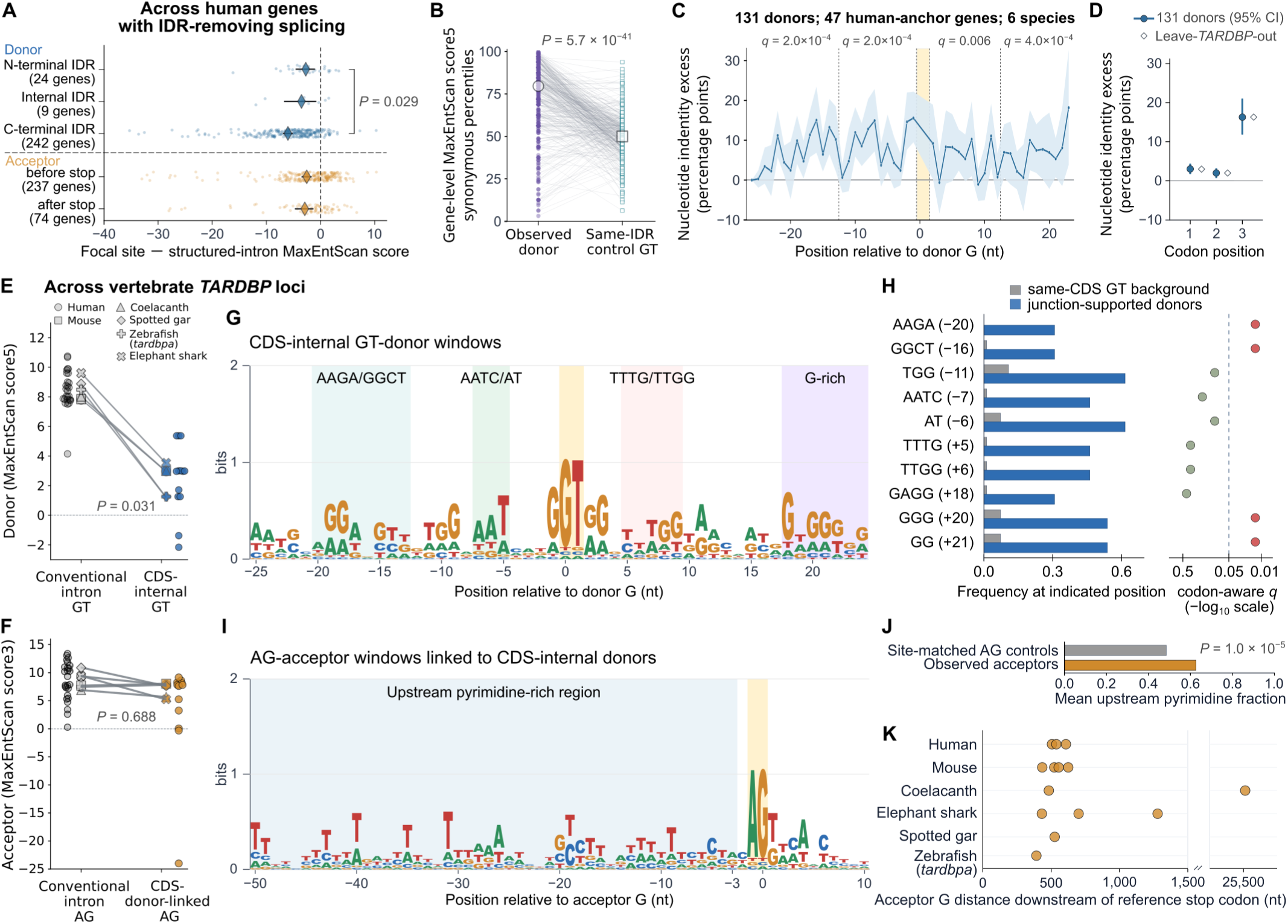
IDR-removing donors have low splice-site scores but rank highly among synonymous alternatives. **(A)** Within-gene differences in median MaxEntScan scores (IDR-removing minus conventional intron sites in annotated structured regions). Donors are grouped by IDR position (N-terminal, internal, C-terminal); acceptors are grouped by position relative to the reference stop codon. Small points represent genes; diamonds and bars show means and bootstrap 95% CIs. The donor-group *P* value is from an omnibus label-permutation test. **(B)** Native 9-nt score5 percentiles among all synonymous alternatives preserving each site’s peptide and central GT. Lines connect within-gene medians for donors and same-IDR control GTs (380 genes); large symbols show across-gene medians. **(C)** Nucleotide identity excess (observed identity to human minus synonymous expectation; percentage points) for 131 non-human donor observations mapped to 47 human genes. Synonymous alternatives preserve each copy’s amino-acid sequence and donor bases and are weighted by CDS codon usage. Displayed *q* values are from gene-level sign-flip tests of four flanking intervals, with Benjamini–Hochberg (BH) adjustment. **(D)** Identity excess by codon position; open diamonds show estimates excluding *TARDBP*. In **C** and **D**, estimates weight genes equally; ribbons or bars show gene-bootstrap 95% CIs. **(E, F)** MaxEntScan score5 for GT donors **(E)** and score3 for linked AG acceptors **(F)**, comparing conventional and CDS-internal splicing across six *TARDBP* loci. Small points represent sites; large symbols show locus medians. *P* values in **B**, **E**, and **F** are from exact two-sided paired sign tests. **(G)** Sequence logo for 13 CDS-internal GT donors (−25 to +24 relative to donor G). **(H)** Frequencies of ten manually selected representative k-mer–position pairs in 13 donor windows and 84 same-species, same-phase coding-region GT controls. Displayed *q* values are from one-sided Fisher exact tests against pooled amino-acid-preserving synonymous-background counts, with BH adjustment across the ten pairs. **(I)** Sequence logo for 14 linked AG acceptors. Blue and yellow shading mark the pyrimidine-rich region (−50 to −3 relative to acceptor G) and AG, respectively. **(J)** Mean C/T fraction in the shaded upstream interval for acceptors versus site-matched post-stop AG controls. *P* is from a one-sided site-stratified permutation test. **(K)** Distance from the reference stop codon to acceptor G; the axis break accommodates the distal coelacanth site. Figure 3 alt text: Plots and sequence logos show low splice-site scores, elevated ranks among synonymous alternatives, and donor-flanking conservation exceeding synonymous expectations for splicing that removes disordered regions.

To account for the amino acids encoded at donor sites, we evaluated donor scores relative to the synonymous sequences available at each coding position. For each eligible donor and same-IDR control GT site, we ranked the native 9-nt sequence by score5 among all synonymous alternatives preserving that site’s local peptide and central GT. Across 380 genes with eligible controls, the median of the within-gene median percentiles was 79.60 for observed donors and 49.93 for controls. The median paired donor-minus-control difference was +27.54 percentile points (95% bootstrap CI, +24.48 to +31.25; exact two-sided sign test, *P* = 5.72 × 10⁻⁴¹), with higher donor ranks in 313 genes, lower ranks in 63, and ties in four (Fig. 3B). Thus, low scores relative to conventional intron donors coexisted with high ranks among the synonymous sequences available at the same coding positions.

In cross-species comparisons, we asked whether donor-flanking nucleotide identity to human exceeded expectations from synonymous alternatives preserving each non-human copy’s local amino-acid sequence. Across 131 non-human donor observations corresponding to 47 human genes, observed identity exceeded these expectations in all four prespecified flanking intervals after excluding donor-constrained positions (BH-adjusted *q* ≤ 0.006; Fig. 3C). The largest excess occurred at third codon positions: observed identity was 55.28%, compared with an expectation of 39.04% (difference, +16.24 percentage points; 95% bootstrap CI, +11.82 to +20.96; BH-adjusted *q* = 1.5 × 10⁻⁴; Fig. 3D). Excluding *TARDBP* yielded nearly identical estimates.

We next asked whether the low donor scores in the human discovery set were also evident across six vertebrate *TARDBP* loci (human, mouse, coelacanth, spotted gar, elephant shark, and zebrafish *tardbpa*). At every locus, the median score5 for CDS-internal GT donors was lower than that for conventional intron donors (13 and 27 sites in total; exact two-sided sign test, unadjusted *P* = 0.031; Fig. 3E). By contrast, median score3 values for linked AG acceptors were lower than those for conventional intron acceptors at four loci and higher at two (14 and 26 sites in total; exact two-sided sign test, unadjusted *P* = 0.688; Fig. 3F). Thus, the reduction in donor scores, but not acceptor scores, was consistent across the six loci.

To characterize donor-flanking sequence features beyond the 9-nt window scored by MaxEntScan, we extended the *TARDBP* donor analysis to 50-nt windows (−25 to +24 relative to donor G). The combined profile of the 13 CDS-internal GT donors showed upstream A/G-rich and AATC/AT-rich regions and downstream T/G-rich and distal G-rich regions (Fig. 3G). The ten sequence–position pairs selected from the exploratory screen were enriched relative to 84 same-species, same-phase control GT sites in the corresponding terminal coding regions. Among the ten selected pairs, four showed enrichment relative to the synonymous background: AAGA at −20, GGCT at −16, and the overlapping GGG and GG k-mers at +20 and +21, respectively (BH-adjusted *q* = 0.0134 for each; Fig. 3H).

We also characterized the 14 linked AG acceptors, which showed a broad upstream pyrimidine-rich profile (Fig. 3I). Their mean C/T fraction from −50 to −3 relative to the acceptor G was 0.626, compared with 0.485 for site-matched post-stop AG controls (one-sided empirical *P* = 1.0 × 10⁻⁵; Fig. 3J). Acceptor positions ranged from several hundred nucleotides to tens of kilobases downstream of the reference stop codon (Fig. 3K). Thus, the sampled *TARDBP* loci combined low donor scores with enriched donor-flanking sequence patterns and pyrimidine-rich acceptor regions at variable downstream distances.

### Teleost ohnologs encode disordered tails but differ in CDS-internal donor usage

We next focused on the splicing contrast between zebrafish *tardbpa* and *tardbpb* observed in the vertebrate survey (Fig. 1F). In both zebrafish and medaka, CDS-internal splicing was prominent at *tardbpa* but weakly supported or undetected at *tardbpb*, whereas post-stop splicing was detected in both copies (Fig. 4A). Both genes were detected across sample–cluster groups in a separate zebrafish adult-brain single-cell RNA-seq dataset (Supplementary Fig. 3A). To determine whether this splicing asymmetry extended more broadly across teleosts, we expanded the comparison to *tardbpa*/*tardbpb* pairs from 15 species, with copy assignments supported by local synteny (Supplementary Table 2). In every pair, IUPred3 predicted disorder-enriched post-RRM2 regions in the IDR-retaining reference translations of both copies (Supplementary Fig. 3B).

**Figure 4.**
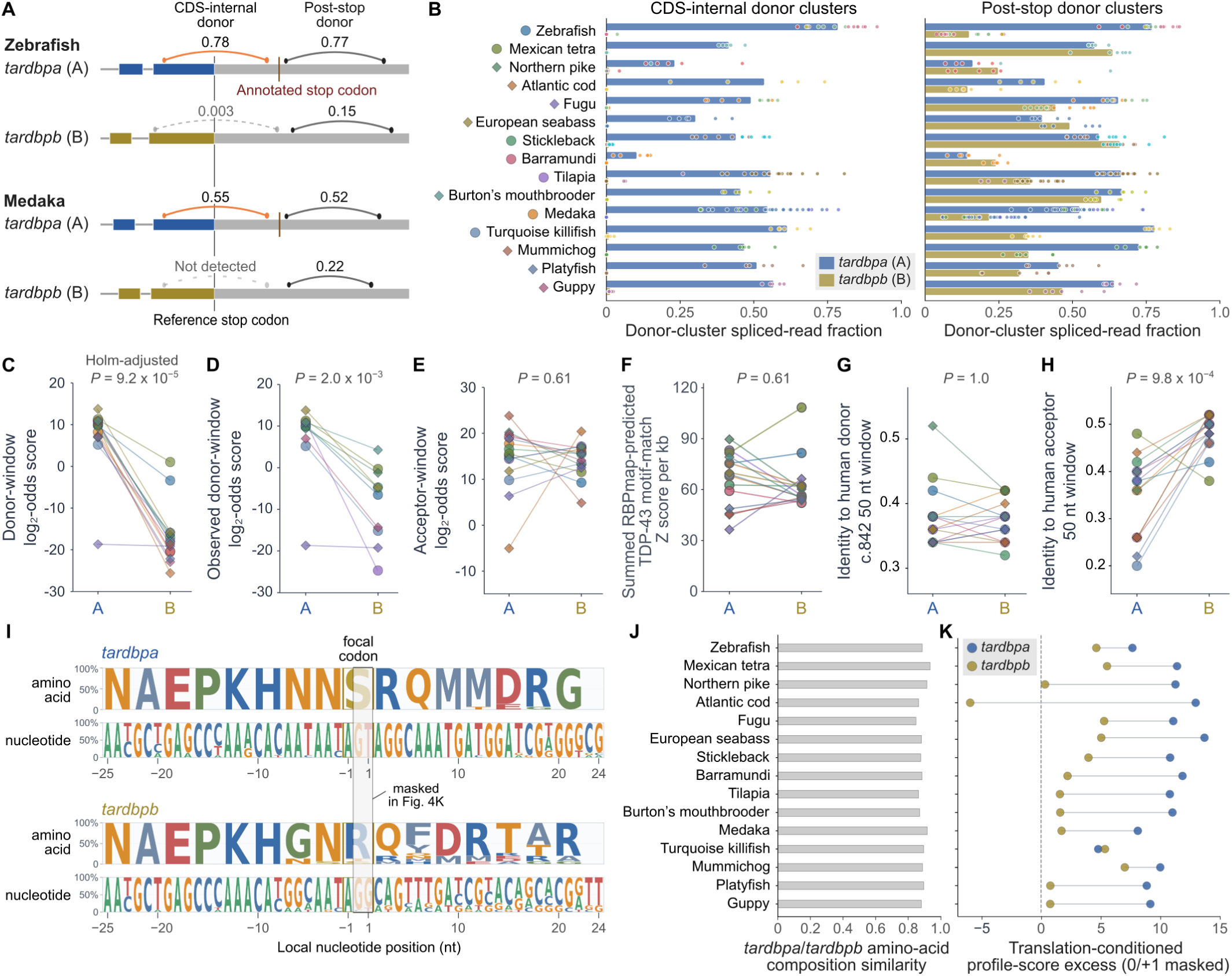
Teleost *tardbpa*/*tardbpb* ohnologs encode compositionally similar post-RRM2 regions but differ in CDS-internal splicing and donor-flanking sequence profiles. **(A)** Zebrafish and medaka terminal regions. Arcs show selected CDS-internal (orange) and post-stop (black) donor clusters; labels indicate locus-level mean spliced-read fractions. Dashed arcs denote low support or no eligible cluster; vertical lines mark stop codons. **(B)** Donor-cluster spliced-read fractions across 15 ohnolog pairs. Points represent RNA-seq runs; bars show within-locus means. Bar and point colors denote copies and projects, respectively. Fractions were not assigned to loci without eligible clusters. **(C)** 50-nt donor-window log₂-odds scores against a 12-site non-teleost profile: observed *tardbpa* donors versus codon-projected *tardbpb* positions. **(D)** Scores for observed GT donors in both copies (ten pairs). **(E)** Acceptor-window log₂-odds scores against a 13-site non-teleost profile: observed *tardbpa* acceptors versus sequence-matched AG candidates in *tardbpb*. **(F)** Summed RBPmap-predicted TDP-43 motif-match Z scores per kb in the 2-kb post-stop region. **(G, H)** Nucleotide identity to human in 50-nt windows around projected human coordinates: c.842 donor **(G)** and acceptor 538 nt downstream of the reference stop **(H)**. **(I)** Amino-acid and nucleotide frequencies in 50-nt windows centered 85 nt downstream of the coding position corresponding to the start of human exon 6 in each copy. In *tardbpa*, centers coincide with RNA-seq-supported donors. Letter height indicates frequency across 15 loci per copy; gold and black boxes mark the focal codon and central dinucleotide, respectively. **(J)** Post-RRM2 amino-acid composition similarity (one minus total variation distance). **(K)** Donor-profile score excess for the windows in **I**, excluding positions 0 and +1: native score minus the median score of 50,000 synonymous-codon permutations preserving each copy’s amino-acid sequence and codon counts. Species order matches **J**. Lines connect species-matched ohnologs in **C– H** and **K**; colors and symbols identify species in **C–H**. CDS-internal cluster detection used an exact one-sided binomial test on discordant pairs. Quantitative comparisons used exact paired sign tests: one-sided for CDS-internal fractions (ten pairs; **B**), **C**, and **K**; two-sided for post-stop fractions **(B)** and **D–H**. All one-sided tests used the alternative *tardbpa* > *tardbpb*. Holm adjustment covers the three primary tests in **B** and **C**; other *P* values are unadjusted (**K**, *P* = 4.88 × 10⁻⁴). Figure 4 alt text: Paired plots and sequence logos show more frequent coding-region donor detection and higher donor-flanking profile scores in tardbpa than tardbpb across fifteen teleost gene pairs.

For quantitative comparisons, eligible junctions sharing the same donor were grouped into clusters, and the cluster with the greatest pooled junction-read support was selected for each locus and donor class. An eligible CDS-internal donor cluster was detected at all 15 *tardbpa* loci and 10 *tardbpb* loci; all five discordant pairs were *tardbpa*-only (exact one-sided binomial test on discordant pairs, Holm-adjusted *P* = 0.031; Fig. 4B). The five *tardbpb* loci without an eligible cluster had complete per-base coverage of their retained coding intervals in all corresponding RNA-seq runs (54 runs in total; Supplementary Table 4). Among the ten pairs with eligible clusters at both loci, donor-cluster spliced-read fractions, averaged across runs within each locus, were higher for *tardbpa* in every pair (across-locus means, 0.485 versus 0.005; exact one-sided paired sign test, Holm-adjusted *P* = 0.00195). In contrast, post-stop donor-cluster spliced-read fractions were higher for *tardbpa* in 10 of 15 pairs (exact two-sided paired sign test, unadjusted *P* = 0.302), and post-stop junctions were detected at every *tardbpb* locus.

Thus, the consistent *tardbpa* bias concerned CDS-internal donor usage, whereas post-stop splicing remained detectable in both ohnologs.

### Donor-flanking profile scores, but not MaxEntScan scores or identity at projected human coordinates, consistently favor *tardbpa*

To identify sequence features associated with the CDS-internal splicing asymmetry, we compared 50-nt windows centered on RNA-seq-supported *tardbpa* donors with windows at codon-projected positions in *tardbpb*. Scores against a profile constructed from 12 non-teleost *TARDBP* CDS-internal GT donors were higher for *tardbpa* in all 15 pairs (mean paired difference, +24.37 log₂-odds; exact one-sided paired sign test, Holm-adjusted *P* = 9.16 × 10⁻⁵; Fig. 4C). Scores were also higher for *tardbpa* in all ten pairs when both windows were centered on observed GT donors (exact two-sided paired sign test, unadjusted *P* = 1.95 × 10⁻³; Fig. 4D and Supplementary Fig. 4). In fugu, however, the predominant *tardbpa* donor, aligned to human c.842, had a relatively low profile score, whereas the upstream donor favored in other teleosts showed substantial use and a high score comparable to those at other *tardbpa* loci. MaxEntScan score5 did not show the same consistent difference when observed *tardbpa* donors were compared with the highest-scoring canonical GT candidates in the predefined *tardbpb* search intervals (Supplementary Fig. 5).

For acceptors, we compared windows around observed *tardbpa* acceptors with sequence-matched AG-centered candidate windows in predefined *tardbpb* search intervals; these candidates were not required to have RNA-seq support. Neither acceptor-profile scores nor summed RBPmap-predicted TDP-43 motif-match Z scores per kilobase in the 2-kb post-stop region showed a consistent directional difference between ohnologs (*tardbpa* > *tardbpb* in 9 of 15 pairs for each; exact two-sided paired sign tests, unadjusted *P* = 0.607 for each; Fig. 4E,F and Supplementary Fig. 6).

We separately compared nucleotide identity to human in 50-nt windows centered on projected human splice-site coordinates. Identity around the c.842 donor position was higher in *tardbpa* in seven pairs and in *tardbpb* in six, with two ties (exact two-sided paired sign test, unadjusted *P* = 1.00; Fig. 4G). By contrast, identity around the major human acceptor, 538 nt downstream of the human reference stop codon, was higher in *tardbpb* in 14 of 15 pairs (mean *tardbpa*-minus-*tardbpb* difference, −13.47 percentage points; exact two-sided paired sign test, unadjusted *P* = 9.77 × 10⁻⁴; Fig. 4H and Supplementary Fig. 7). Thus, nucleotide identity around the projected human splice-site coordinates did not show the consistent *tardbpa* bias observed for donor-profile scores.

We next asked whether the donor-profile difference persisted after excluding the central dinucleotide from scoring and accounting for the encoded amino-acid sequences. For this analysis, we used 50-nt windows centered 85 nt downstream of the coding position corresponding to the start of human exon 6 in each copy. At this offset, every *tardbpa* locus contained an AGT (Ser) codon whose final two bases formed an RNA-seq-supported GT donor, whereas most *tardbpb* loci carried AGG (Arg) (Fig. 4I). Despite residue-level divergence (median residue identity, 0.634), the paired post-RRM2 regions had similar amino-acid compositions (median composition similarity, 0.885; Fig. 4J). We excluded positions 0 and +1 from scoring and compared native scores with scores from 50,000 synonymous-codon permutations for each copy, preserving the encoded amino-acid sequence and codon counts. Donor-profile score excess, defined as the native score minus the median permuted-sequence score, was higher in *tardbpa* in 14 of 15 pairs (median paired difference, +8.10 log₂-odds; exact one-sided paired sign test, unadjusted *P* = 4.88 × 10⁻⁴; Fig. 4K). Together, these comparisons associate the CDS-internal splicing asymmetry with a donor-flanking profile difference that extends beyond the central dinucleotide and persists relative to copy-specific synonymous expectations.

### Donor-proximal *cis* sequences regulate donor choice and splice-product balance

To test whether donor-proximal sequence differences altered splicing, we next introduced reciprocal substitutions into zebrafish *tardbpa* and *tardbpb* minigenes. We exchanged the central dinucleotide alone, the dinucleotide together with the downstream G-rich sequence, or broader intervals spanning −25 to +24 or −40 to +40 nt (Fig. 5A,E). Minigenes were assayed in HEK293T cells under basal TDP-43 conditions. RT-qPCR quantified *In* relative to the constitutive-junction signal (*In*/*CJ*), and endpoint RT-PCR resolved splice products and measured *Ex*/*In* band-intensity ratios (Fig. 5A,E).

**Figure 5.**
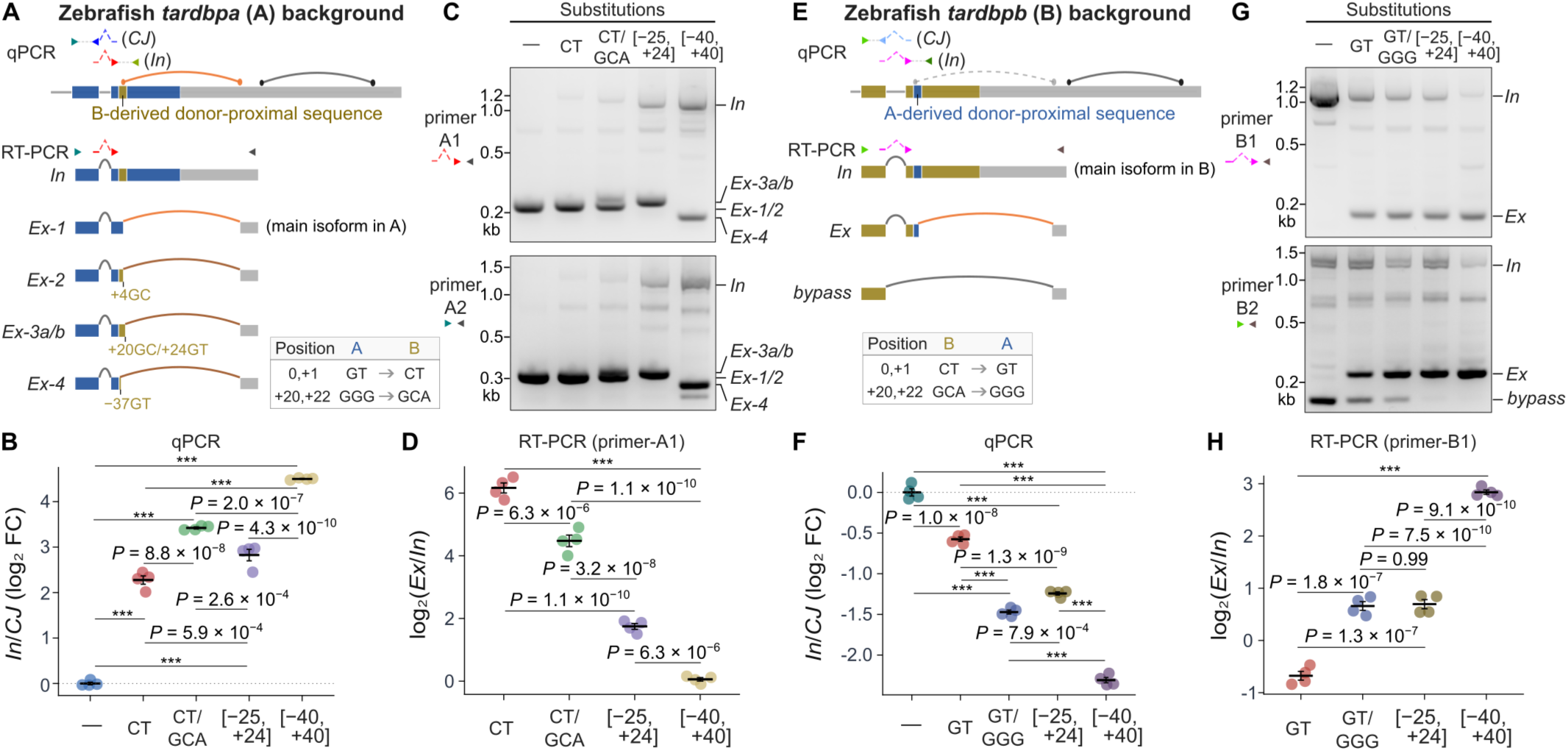
Donor-proximal sequence substitutions alter donor choice and splice-product balance. **(A, E)** Schematics of reciprocal substitutions in zebrafish *tardbpa* **(A)** and *tardbpb* **(E)** minigenes, assayed in HEK293T cells under basal TDP-43 conditions. Coordinates refer to the *tardbpa* reference donor and its aligned position in *tardbpb*; bracketed intervals were replaced with aligned sequence from the other ohnolog. *In* denotes the IDR-retaining product; *Ex*-1 **(A)** and *Ex* **(E)** are IDR-removing products using the reference CDS-internal donor. *Ex*-2, *Ex*-3a, *Ex*-3b, and *Ex*-4 use donors at +4 (GC), +20 (GC), +24 (GT), and −37 (GT), respectively. *Ex*-3a and *Ex*-3b are unresolved on gels. The *bypass* product joins the upstream constitutive-intron donor to a downstream 3′UTR acceptor. Arrowheads indicate primer positions. **(B, F)** RT-qPCR quantification of the *In* signal normalized to the constitutive-junction (*CJ*) signal in *tardbpa* **(B)** and *tardbpb* **(F)** constructs. The *CJ* assay detects transcripts containing the exon–exon junction formed by removal of the upstream constitutive intron and excludes bypass transcripts. *In*/*CJ* ratios are expressed as log₂ fold change relative to the mean *In*/*CJ* value of the corresponding unsubstituted control group (−). **(C, G)** Endpoint RT-PCR using primer sets A1 **(C, top)**, A2 **(C, bottom)**, B1 **(G, top)**, and B2 **(G, bottom)**. A1/B1 forward primers span the constitutive exon–exon junction; A2/B2 forward primers lie within the upstream exon, allowing *bypass* detection. **(D, H)** *Ex*/*In* band-intensity ratios from A1 **(D)** and B1 **(H)**, expressed as log₂(*Ex*/*In*). *Ex* is the summed intensity of the indicated IDR-removing bands, excluding *bypass*. Unsubstituted controls were excluded from *Ex*/*In* quantification and associated tests because *In* (*tardbpa*) or *Ex* (*tardbpb*) was undetectable by endpoint RT-PCR. Horizontal bars and error bars show mean ± SEM (*n* = 4). *P* values are Tukey-adjusted following one-way ANOVA (\*\*\**P* < 1.0 × 10⁻¹⁰). Figure 5 alt text: Schematics, gels, and quantitative plots show that reciprocal donor-proximal substitutions alter donor choice and the balance of inclusion, exclusion, and bypass products in zebrafish minigenes.

The unsubstituted *tardbpa* minigene predominantly produced *Ex-1*, the IDR-removing transcript using the reference GT donor (Fig. 5B–D). Replacing the reference GT with the CT present at the corresponding *tardbpb* position increased *In*/*CJ* but did not abolish CDS-internal splicing; instead, use of a GC donor at +4 generated *Ex*-2. Replacing the downstream GGG triplet with GCA reduced *Ex*/*In* relative to CT alone. Broader *tardbpb*-derived substitutions also lowered *Ex*/*In*, with the −40 to +40 construct yielding the lowest *Ex*/*In* and highest *In*/*CJ* among the substituted constructs. These broader replacements also altered donor choice, generating *Ex*-3a and *Ex*-3b from donors at +20 (GC) and +24 (GT), respectively, and *Ex*-4 from a donor at −37 (GT) (Fig. 5C; Supplementary Table 6). Thus, disrupting the predominant GT donor redirected splicing to alternative donors rather than eliminating IDR-removing products.

In the *tardbpb* background, the unsubstituted minigene predominantly produced *In* and also yielded a *bypass* product joining the donor of the upstream constitutive intron to a downstream 3′UTR acceptor (Fig. 5E–H). This *bypass* junction was also detected with low read support in native zebrafish *tardbpb* RNA-seq data, although the corresponding *bypass* product was substantially more prominent in the minigene assay. Introducing GT at the reference position generated *Ex*, but *In* remained the major product. Adding the downstream GCA-to-GGG substitution increased *Ex*/*In* relative to GT alone. The −25 to +24 substitution yielded a mean *Ex*/*In* similar to that of the GT-plus-GGG construct, whereas the −40 to +40 substitution gave the highest *Ex*/*In* and lowest *In*/*CJ* among the substituted constructs. RT-PCR using an upstream-exon primer also showed a shift toward *Ex* relative to the *bypass* product (Fig. 5G).

The *bypass* product showed that the downstream acceptor was already used in the unsubstituted *tardbpb* minigene. With this acceptor region unchanged, transferring *tardbpa* donor-proximal sequence shifted product balance toward use of the CDS-internal donor. Together, the reciprocal substitutions establish donor-proximal *cis* control beyond the central GT, affecting alternative donor choice and the balance of IDR-retaining and IDR-removing splice products in these minigenes.

### Donor-proximal substitutions alter reporter-product balance and puncta formation

To determine whether the changes in splice-product balance were reflected at the protein level, we next analyzed EGFP-tagged versions of the same zebrafish minigenes in HEK293T cells by immunoblotting and live-cell imaging (Fig. 6A,J). GFP-reactive bands in the RIPA-soluble fraction were assigned operationally from the corresponding RNA products, apparent molecular masses, and responses to *In*-selective siRNA (Fig. 6A–D,J–M and Supplementary Table 6).

**Figure 6.**
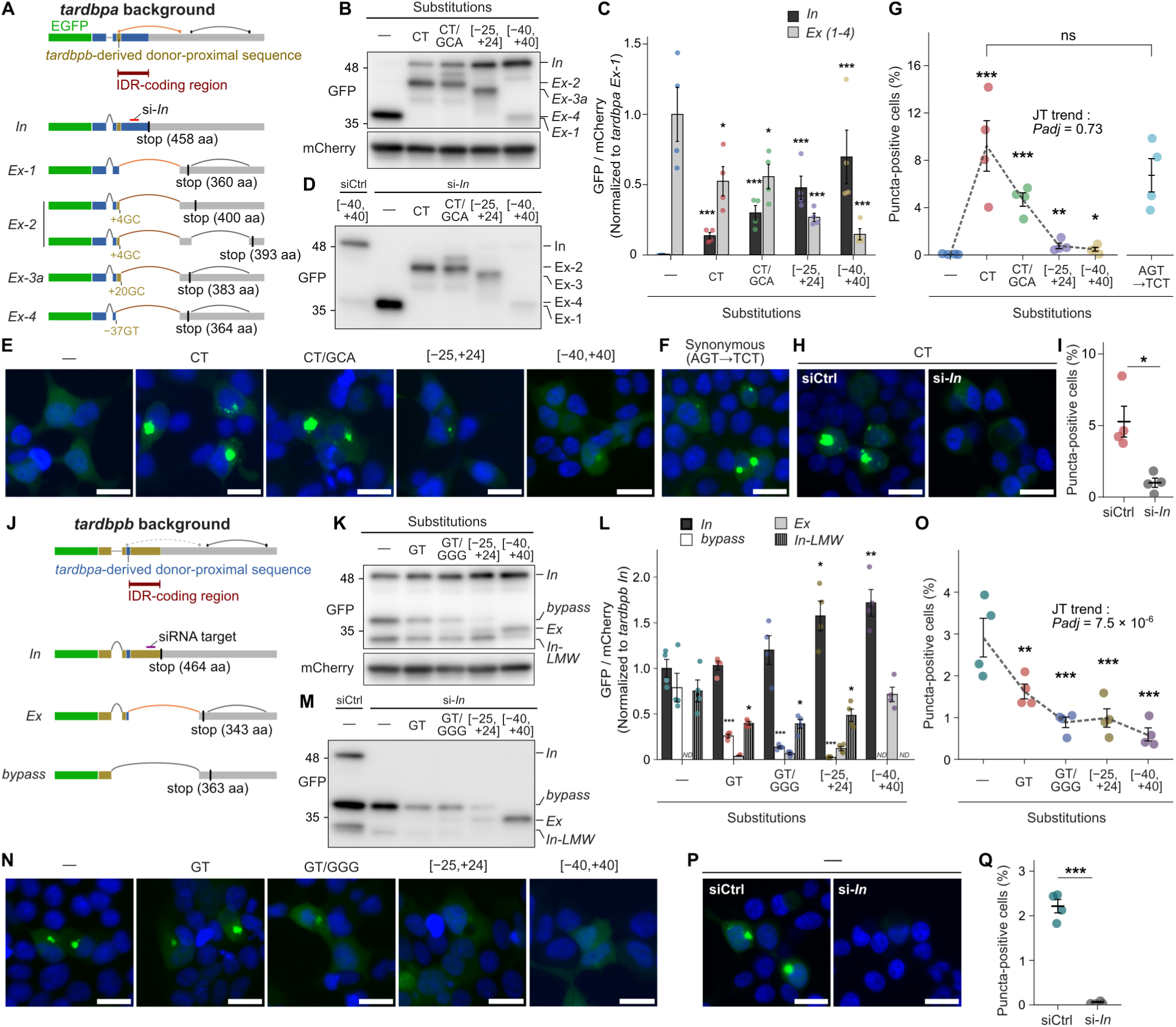
Donor-proximal substitutions alter reporter-product balance and puncta formation. **(A, J)** Schematics of EGFP-tagged zebrafish *tardbpa* **(A)** and *tardbpb* **(J)** terminal-region reporters assayed in HEK293T cells and representative products. *In* denotes the IDR-retaining reporter product; *Ex* denotes IDR-removing products. *Bypass* joins the upstream constitutive-intron donor to a downstream 3′UTR acceptor. *In-LMW* denotes an *In*-associated lower-molecular-weight species. Bands were assigned operationally from RNA products, apparent molecular masses, and siRNA responses. −, unsubstituted reporter. **(B, K)** Anti-GFP and anti-mCherry immunoblots of RIPA-soluble fractions from the *tardbpa* **(B)** and *tardbpb* **(K)** substitution series. **(C, L)** Quantification of *In* and summed *Ex* products **(C)**, or *In*, *bypass*, *Ex*, and *In-LMW* **(L)**, normalized to mCherry and expressed relative to *tardbpa Ex-1* **(C)** or *tardbpb In* **(L)**. ND, not detected. **(D, M)** Anti-GFP immunoblots following *In*-selective siRNA (si-*In*) treatment. Non-targeting siRNA (siCtrl) lanes show the −40 to +40 *tardbpa* reporter **(D)** and parental *tardbpb* reporter **(M)**. **(E, F, N)** Representative fluorescence images of cells expressing the indicated *tardbpa* **(E)**, synonymous AGT→TCT **(F)**, and *tardbpb* **(N)** reporters, with puncta quantification in **G** and **O**. **(H, P)** Images of the *tardbpa* CT reporter **(H)** and parental *tardbpb* reporter **(P)** with siCtrl or si-*In*, quantified in **I** and **Q**. Green, EGFP; blue, Hoechst-stained nuclei. Scale bars, 20 µm. Puncta-positive cell fractions were calculated among mCherry-positive cells. Points represent culture wells (*n* = 4 per condition); data are presented as mean ± SEM. In **C** and **L**, comparisons with the parental reporter used ANOVA-based contrasts with Holm adjustment. Values for the *In* product in **C** were log₂-transformed. *Ex* abundance in **L** was not tested against the parental value because no parental *Ex* band was detected. In **G** and **O**, comparisons with the parental reporter used two-sided Dunnett-type contrasts from beta-binomial generalized linear mixed-effects models with single-step multivariate-*t* adjustment. TCT was compared separately with CT. One-sided Jonckheere–Terpstra tests assessed ordered substitution trends, with Holm adjustment across the two backgrounds; TCT was excluded. **I** and **Q** used two-sided Welch’s *t*-tests. \**P* < 0.05, \*\**P* < 0.01, \*\*\**P* < 0.001; ns, not significant. Adjusted *P* values are shown where applicable. Figure 6 alt text: Immunoblots and microscopy show altered protein-product balance and puncta after donor-proximal substitutions in terminal-region reporters in HEK293T cells; targeted depletion of inclusion transcripts reduces puncta.

In the *tardbpa* background, the unsubstituted reporter predominantly produced the short *Ex-1* product and showed diffuse EGFP fluorescence (Fig. 6B,C,E). The GT-to-CT substitution increased *In*-product abundance and the fraction of puncta-positive cells (Fig. 6B,C,E,G). A synonymous AGT→TCT substitution that also disrupted the donor while retaining serine yielded RNA and reporter-product patterns similar to those seen with CT and a similar punctate phenotype (Supplementary Fig. 8A,B; Fig. 6F,G). *In*-selective siRNA depleted the *In* product in the *tardbpa* reporter assay (Fig. 6D) and markedly reduced puncta in cells expressing the CT reporter (Fig. 6H,I). These results support a contribution of altered RNA processing to puncta formation without requiring the amino-acid change introduced by the CT substitution. Across the substitution series, however, the mean puncta-positive fraction peaked with CT and was lower with broader replacements, despite their higher *In*/*CJ* RNA ratios and soluble *In*-product abundance (Figs. 5B and 6B,C,E,G).

The unsubstituted *tardbpb* reporter produced *In* and *bypass* products together with an *In*-associated lower-molecular-weight species (*In-LMW*); cells expressing this reporter formed cytoplasmic puncta (Fig. 6K,L,N). *In*-selective siRNA reduced both *In* and *In-LMW* and nearly abolished puncta, while the *bypass* product remained detectable (Fig. 6M,P,Q). Introducing *tardbpa*-derived donor-proximal sequences increased *Ex*-product abundance and reduced *bypass*-product abundance. *In-LMW* was also reduced and was undetectable in the −40 to +40 construct (Fig. 6K,L). Puncta-positive fractions showed an overall decreasing trend across the ordered substitutions (Holm-adjusted Jonckheere–Terpstra *P* = 7.5 × 10⁻⁶; Fig. 6N,O), even though soluble *In*-product abundance was not decreased (Fig. 6K,L).

Together, these experiments link donor-proximal regulation of splicing to reporter-product balance and puncta formation. Puncta formation was reduced by siRNA targeting the IDR-retaining transcript in both backgrounds, but did not consistently track the abundance of the soluble long *In* product.

## Discussion

This study shows that exitron-type splicing predicted to remove IDRs can recur across jawed vertebrates despite divergence in coding sequence and variation in splice-junction positions. The shared predicted outcome is loss of most of the targeted IDR residues while retaining all annotated structured-region residues unchanged. Because IDRs mediate functional interactions but can also promote aberrant self-association [5,6], regulating their inclusion offers a potential means of modulating these properties through RNA processing.

Splicing-associated nucleotide features and flexibility in splice-site placement may help explain how divergent coding sequences yield similar predicted IDR-removing outcomes. Comparisons with synonymous alternatives encoding the same amino-acid sequence revealed high donor-score ranks and excess donor-flanking nucleotide identity, despite low scores relative to conventional intron donors and modest absolute identity. For C-terminal IDRs, no downstream structured region needs to be reconnected in frame, allowing donor, acceptor, and termination positions to vary while leaving upstream structured-region sequences unchanged.

In *TARDBP*, the sequence encoding the disordered tail also carries *cis* information that regulates IDR retention. In zebrafish minigenes, disrupting the predominant *tardbpa* donor redirected excision to nearby sites, whereas introducing GT into *tardbpb* generated *Ex* while leaving *In* as the major product. Broader flanking-sequence exchanges changed donor choice and splice-product proportions with the downstream acceptor region held constant. The functional relevance of IDR-retaining protein products is supported by earlier zebrafish studies in which increased production of a long *tardbpa* isoform compensated for *tardbpb* loss [61,62]. The donor neighborhood therefore regulates both the choice of junction and the balance between IDR-retaining and IDR-removing transcripts.

This donor-proximal regulation also influenced puncta formation in *TARDBP* terminal-region reporters. A synonymous donor-disrupting substitution shifted the balance of both transcripts and reporter proteins toward IDR-retaining forms and promoted puncta formation, whereas *In*-selective depletion suppressed puncta in both reporter backgrounds. These complementary perturbations support a contribution of RNA processing to the phenotype. Across the broader substitutions, puncta formation did not consistently track the abundance of the soluble long *In* product, and the accompanying amino-acid changes may also contribute to the phenotype. Overall, RNA-level control of IDR retention can influence the cellular behavior of the encoded products.

This study has several limitations. The organization of donor-proximal cis-regulatory sequences and the evolutionary history of IDR-removing splicing remain unresolved. Systematic synonymous mutagenesis at endogenous *TARDBP* loci could resolve the contributions of localized motifs and broader sequence features to splice-site choice and full-length TDP-43 output. Testing these features in native *trans* contexts would assess whether the regulatory activities observed in human cells also operate in their species of origin. Combining these functional comparisons with phylogenetic analyses could help determine whether IDR-removing splicing in *TARDBP* was retained from a common ancestor or acquired independently in different lineages.

Taken together, these findings suggest that variation in IDR-coding sequences can influence protein behavior through changes in the encoded IDR, the regulation of its inclusion in protein products, or both [2,63]. Determining how changes in IDR inclusion affect molecular interactions and self-association in native cellular contexts will clarify the physiological significance of the option to retain or remove an IDR.

## Acknowledgements

Artificial-intelligence tools, including ChatGPT, were used for code drafting, debugging, and English-language editing. All outputs were reviewed by the authors, who take full responsibility for the analyses and manuscript.

## Author contributions

Conceptualization: T.Y. (supporting) and A.S. (lead); Experimental design: T.Y. and A.S.; Computational analyses: A.S.; Experiments: T.Y., S.K., G.T., A.W., and S.O.; Visualization: T.Y. (supporting) and A.S. (lead); Supervision: K.Y. and O.O.; Writing (original draft): T.Y. (supporting) and A.S. (lead); Writing (review and editing): T.Y., S.K., G.T., K.Y., O.O., and A.S.

