## Supplementary_Figures for "Exitron-type splicing of intrinsically disordered regions recurs across jawed vertebrates despite sequence divergence: donor-proximal *cis* control in *TARDBP*"

### Supplementary Figures 1–8

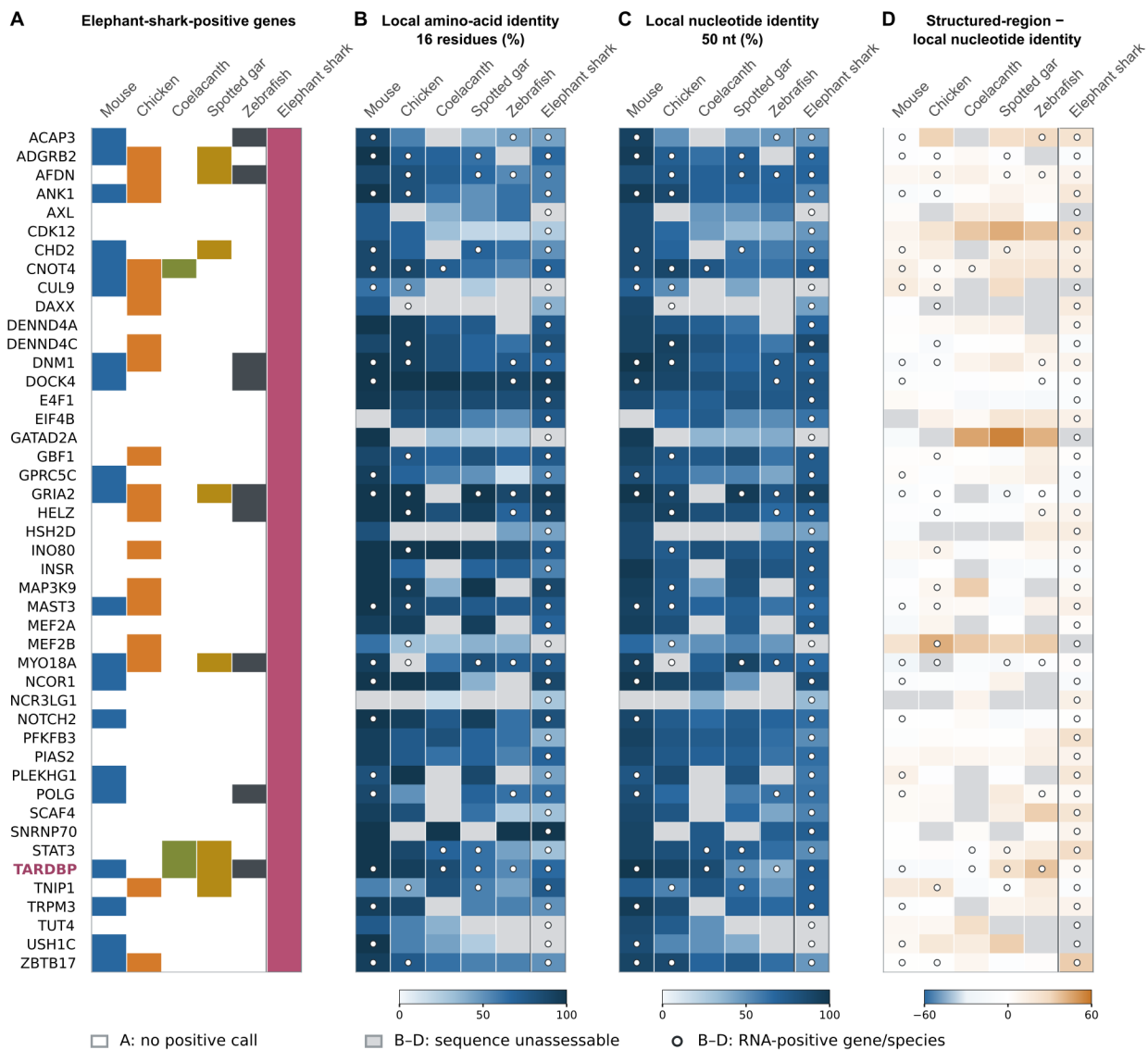

**Supplementary Figure 1. Donor-neighborhood sequence conservation in genes with IDR-removing splicing in elephant shark.** Rows represent the 45 human genes with qualifying splicing in elephant shark (Fig. 1C); columns show six non-human species, with elephant shark outlined. **(A)** Colored cells indicate $\geq 1$ qualifying exact donor-acceptor pair detected in $\geq 2$ physical RNA-seq runs; white cells indicate no qualifying call. **(B, C)** Amino-acid **(B)** and nucleotide **(C)** identity to human in human-donor-centered 50-nt coding windows ($-25$ to $+24$), with matches divided by 16 residues and 50 nt, respectively. Amino-acid comparisons use the 16 complete codons within each window. **(D)** Nucleotide-identity difference (annotated structured regions minus local windows; percentage points); positive values indicate lower local identity. Estimates were averaged across assessable copies per donor, then across assessable donors per gene-species combination. Sequence assessability required $\geq 80\%$ local and structured-region coverage at both sequence levels, yielding 39 assessable elephant shark genes. White circles in B-D mark RNA-positive calls independently of sequence assessability; gray cells indicate missing estimates, including six RNA-positive elephant shark genes.

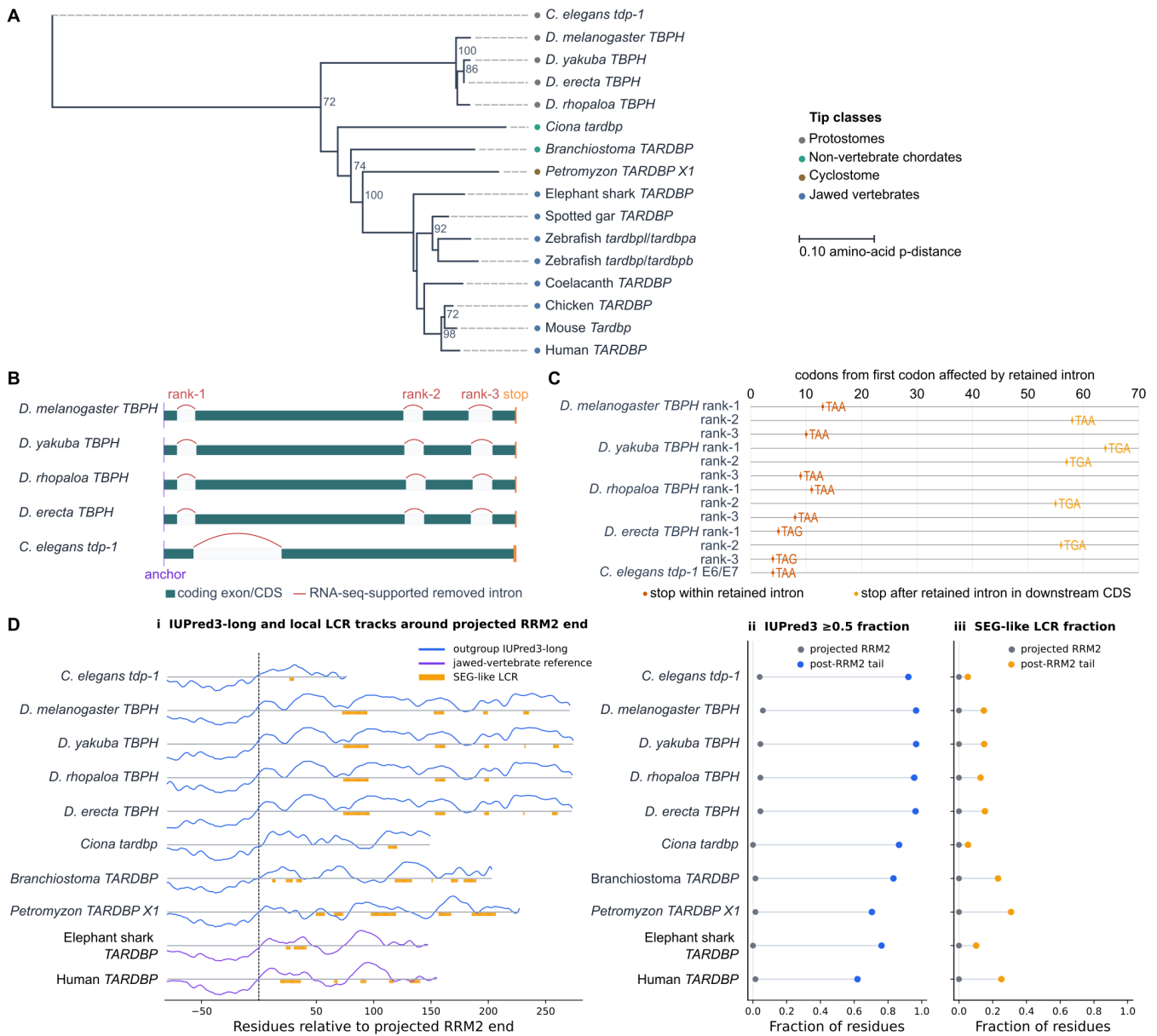

**Supplementary Figure 2. Phylogenetic context, protostome terminal coding-region organization, and post-RRM2 sequence properties in TDP-43-family outgroups. (A)** Structured-core phylogeny used for the outgroup survey, comprising the eight jawed-vertebrate proteins displayed in Fig. 1E and eight outgroup homologs. Regions corresponding to human TDP-43 residues 1–259 were aligned with MAFFT; the alignment was filtered to retain columns with $\leq 50\%$ gaps, and a neighbor-joining tree was inferred from amino-acid p-distances with 200 bootstrap replicates. **(B)** Terminal coding-region organization from the anchor corresponding to the start of human exon 6 to the stop codon in four drosophilid *TBPH* loci and *C. elegans tdp-1*. Teal boxes denote coding segments, orange the stop codon, and red arcs annotated introns supported by RNA-seq junction reads. **(C)** Predicted coding consequence of retaining each protostome terminal-region intron. The position of the first in-frame stop codon is measured in codons from the first affected codon; red indicates a stop within the retained intron, and orange indicates a stop in the downstream coding sequence. **(D)** IUPred3-long disorder and entropy-based low-complexity region (LCR) tracks for eight outgroup homologs, with human and elephant shark references, aligned at the projected RRM2 endpoint (i). Summary plots compare fractions of residues with IUPred3-long scores $\geq 0.5$ (ii) or classified as low complexity (iii) between the post-RRM2 tail and RRM2 reference region.

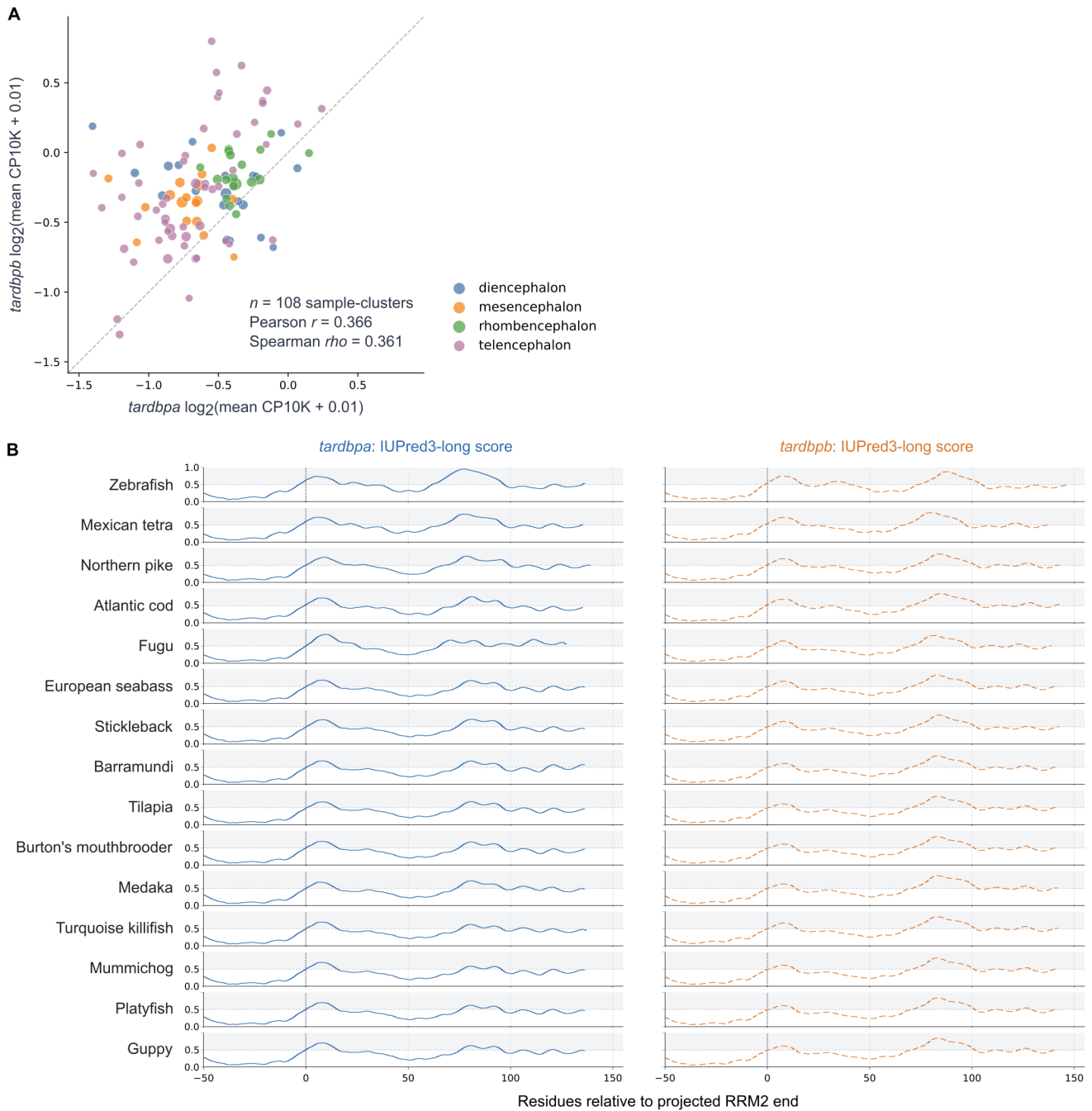

**Supplementary Figure 3. *TARDBP* ohnolog expression in zebrafish brain and predicted post-RRM2 disorder across teleosts. (A)** Expression of zebrafish *tardbpa* (*tardbpl*) and *tardbpb* (*tardbp*) from ten processed GSE246714 adult-brain single-cell RNA-seq loom matrices (48,578 cells). Points represent GEO sample-cluster groups with $\geq 20$ cells ($n = 108$), plotted as $\log_2(\text{mean CP10K} + 0.01)$; CP10K denotes counts per 10,000 UMIs. Point color and size indicate brain region and cell number, respectively; the dashed line denotes equal expression. Supplied numeric cluster labels were not interpreted as annotated cell types. Pearson and Spearman coefficients summarize expression covariation across groups. **(B)** IUPred3-long disorder profiles for IDR-retaining reference translations of 15 teleost *tardbpa/tardbpb* pairs, aligned at the projected RRM2 endpoint (position 0). Blue solid and orange dashed lines show *tardbpa* and *tardbpb*, respectively; horizontal dotted lines mark the 0.5 threshold.

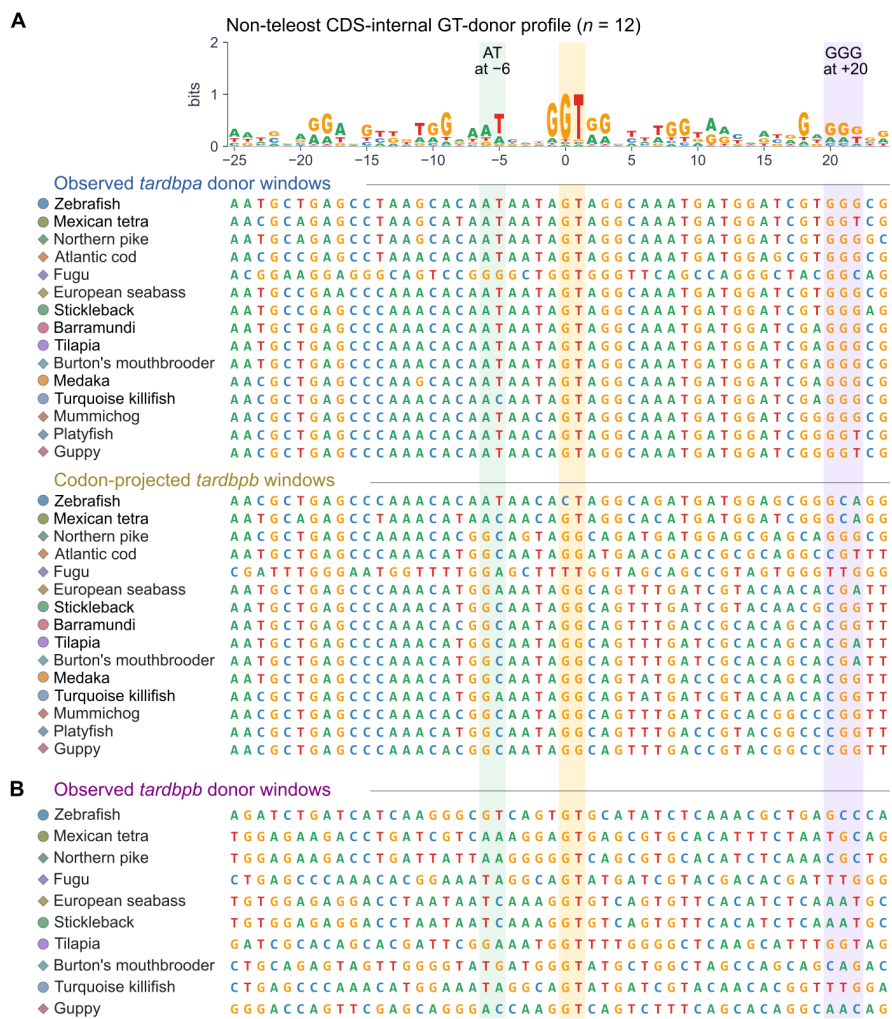

**Supplementary Figure 4. Sequence windows underlying teleost *tardbpa/tardbpb* donor-profile score comparisons.** The reference logo represents 12 non-teleost CDS-internal GT donors, with donor G at position 0. All 50-nt windows (−25 to +24) are shown in transcript orientation. Shaded columns mark the reference-profile positions of AT beginning at −6, the central dinucleotide (0 and +1), and GGG beginning at +20. **(A)** Observed *tardbpa* donor windows (upper block) and codon-projected *tardbpb* windows (lower block) across 15 species. The projected *tardbpb* positions were defined independently of junction detection and need not correspond to observed splice donors. Paired log<sub>2</sub>-odds scores are shown in Fig. 4C. **(B)** Observed *tardbpb* GT donor windows for the ten species with eligible CDS-internal donor clusters, centered on the RNA-seq-supported donor G. These donors may occupy different coding positions from the paired *tardbpa* donors in A; paired scores are shown in Fig. 4D.

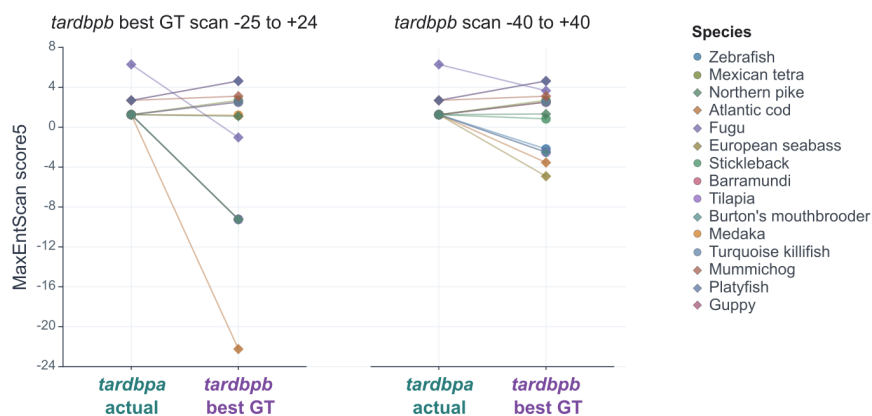

**Supplementary Figure 5. MaxEntScan donor scores do not show the consistent *tardbpa*–*tardbpb* difference observed for donor-profile scores.** MaxEntScan score5 values are shown for RNA-seq-supported *tardbpa* CDS-internal donors and the highest-scoring canonical GT candidate in each paired *tardbpb* search interval. Intervals were centered on the nucleotide-projected *tardbpa* donor position: –25 to +24 nt (left) and –40 to +40 nt (right; sensitivity analysis). Pairs lacking a canonical GT candidate in the relevant interval were excluded. Lines connect species-matched ohnologs; colors and symbols identify species. The *tardbpa* score was higher in 8 of 14 evaluable pairs in the primary scan and 7 of 15 in the sensitivity scan, unlike the uniformly higher *tardbpa* donor-profile scores in Fig. 4C.

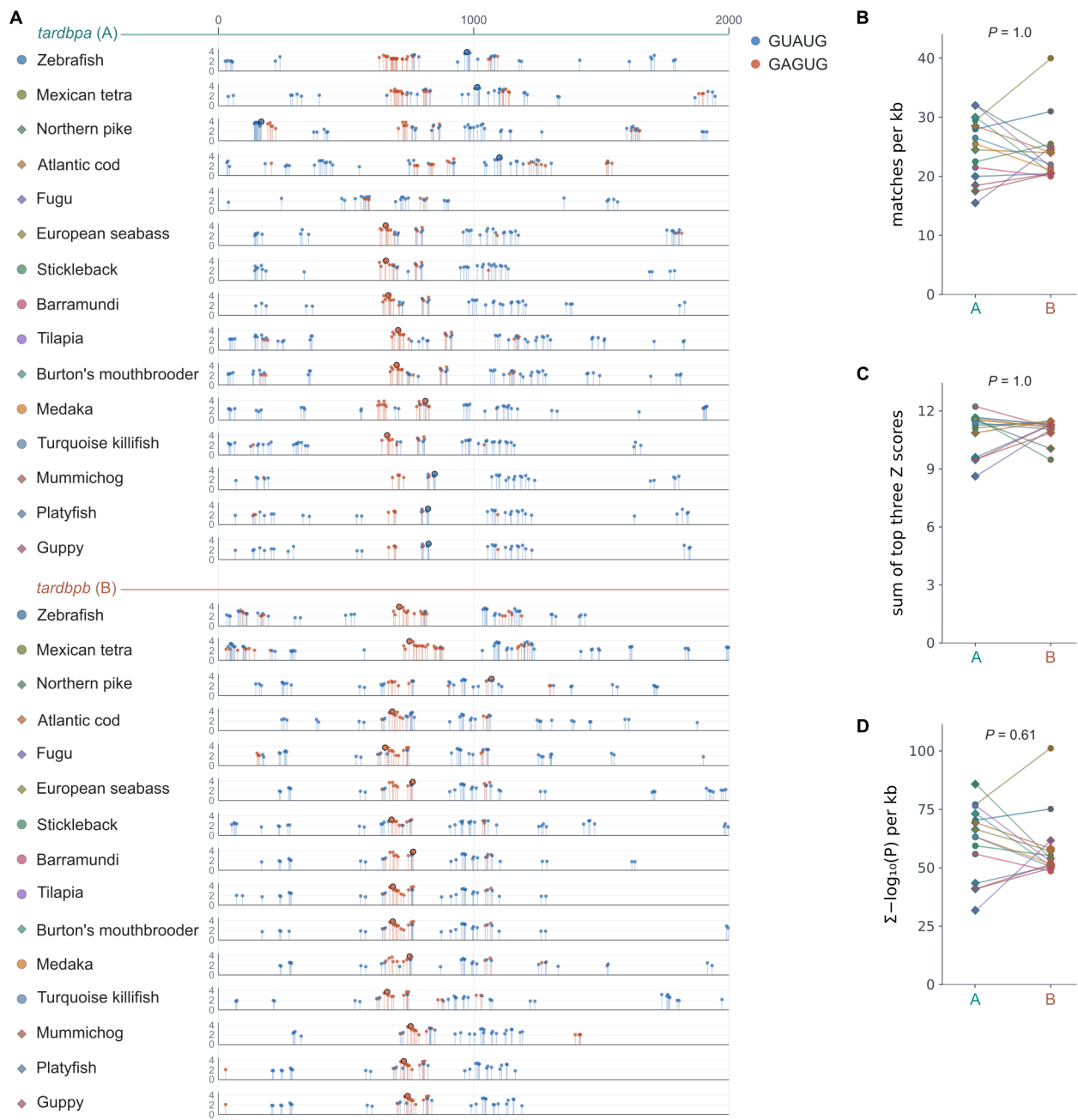

**Supplementary Figure 6. Distribution and alternative summaries of RBPmap-predicted TDP-43 motif matches in teleost post-stop regions. (A)** Matches to *TARDBP* motif models 284 and 285 from +1 to +2,000 nt downstream of the reference stop codon across 15 *tardbpa/tardbpb* pairs. Rows represent loci, grouped into *tardbpa* (upper) and *tardbpb* (lower) blocks. Points show stop-relative positions and RBPmap Z scores. Blue and red-brown denote GUAUG and GAGUG motif classes, respectively; black outlines mark the highest-scoring match per locus. **(B–D)** Paired comparisons of predicted-match density per kilobase **(B)**, the sum of the three highest Z scores **(C)**, and summed $-\log_{10}$-transformed motif-match *P* values per kilobase **(D)**. Lines connect species-matched ohnologs; point colors and symbols identify species. These secondary comparisons used exact two-sided paired sign tests; displayed *P* values are unadjusted.

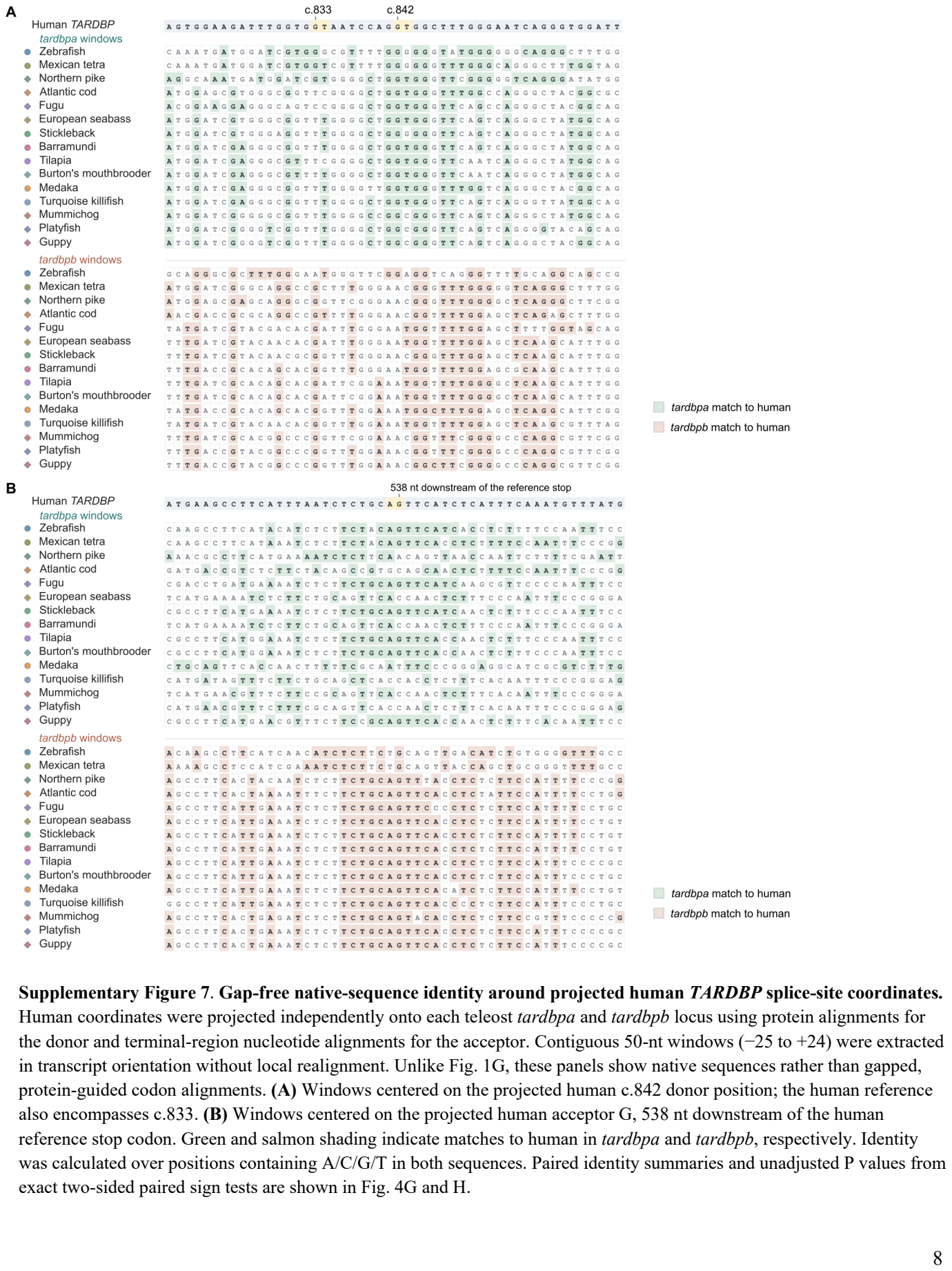

Supplementary Figure 7. Gap-free native-sequence identity around projected human TARDBP splice-site coordinates.

Human coordinates were projected independently onto each teleost tardbpa and tardbpb locus using protein alignments for the donor and terminal-region nucleotide alignments for the acceptor. Contiguous 50-nt windows (−25 to +24) were extracted in transcript orientation without local realignment. Unlike Fig. 1G, these panels show native sequences rather than gapped, protein-guided codon alignments. (A) Windows centered on the projected human c.842 donor position; the human reference also encompasses c.833. (B) Windows centered on the projected human acceptor G, 538 nt downstream of the human reference stop codon. Green and salmon shading indicate matches to human in tardbpa and tardbpb, respectively. Identity was calculated over positions containing A/C/G/T in both sequences. Paired identity summaries and unadjusted P values from exact two-sided paired sign tests are shown in Fig. 4G and H.

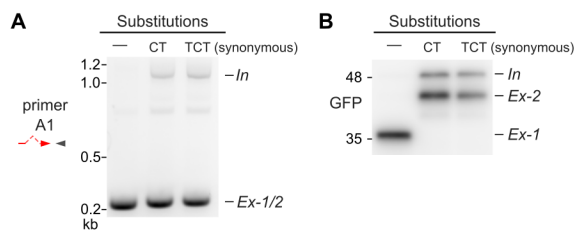

**Supplementary Figure 8. A synonymous donor-disrupting substitution yields splice- and reporter-product patterns similar to those of the CT substitution.** EGFP-tagged zebrafish *tardbpa* reporters carrying no substitution (—), a GT-to-CT substitution (CT), or a synonymous AGT→TCT substitution (TCT) were analyzed in HEK293T cells. TCT disrupts the donor while retaining the encoded serine. **(A)** Endpoint RT-PCR using primer set A1. In denotes the IDR-retaining transcript; *Ex-1/2* denotes unresolved IDR-removing products. **(B)** Anti-GFP immunoblot of the corresponding reporters, with In, *Ex-1*, and *Ex-2* products indicated. Puncta phenotypes are shown in Fig. 6F,G.
